# Oncogenetic Landscape Of Lymphomagenesis In Coeliac Disease

**DOI:** 10.1101/2020.09.07.275032

**Authors:** Sascha Cording, Ludovic Lhermitte, Georgia Malamut, Sofia Berrabah, Amélie Trinquand, Nicolas Guegan, Patrick Villarese, Sophie Kaltenbach, Bertrand Meresse, Sherine Khater, Michael Dussiot, Marc Bras, Morgane Cheminant, Bruno Tesson, Christine Bole-Feysot, Julie Bruneau, Thierry Jo Molina, David Sibon, Elizabeth Macintyre, Olivier Hermine, Christophe Cellier, Vahid Asnafi, Nadine Cerf-Bensussan

**Affiliations:** Université de Paris, Imagine Institute, Laboratory of Intestinal Immunity, INSERM UMR 1163, F-75015, Paris, France; Université de Paris, Institut Necker-Enfants Malades, INSERM UMR 1151, F-75015, Paris, France; Laboratory of Onco-Haematology, AP-HP, Hôpital Necker Enfants-Malades, F-75015, Paris, France; Department of Gastroenterology, AP-HP, Hôpital Européen Georges Pompidou, F-75015, Paris, France; Department of Gastroenterology, AP-HP, Hôpital Cochin, F-75015, Paris, France; National Children’s Research Centre, Children’s Health Ireland at Crumlin, Dublin, Ireland; Department of Cytogenetics, AP-HP, Hôpital Necker-Enfants Malades, F-75015, Paris, France; Université de Lille, CHU Lille, INSERM UMR 1286 – IFNINITE – Institute for Translational Research in Inflammation, F-59000 Lille, France; Université de Paris, Imagine Institute, Laboratory of molecular mechanisms of hematological disorders and therapeutic implications, INSERM UMR 1163, F-75015, Paris, France; Université de Paris, Institut Imagine, Bioinformatics Platform, F-75015, Paris, France; Institut Carnot CALYM, F-75015 Paris, France; Université de Paris, Imagine Institute, Genomics Platform, F-75015, Paris, France; Department of Pathology, AP-HP, Hôpital Necker-Enfants Malades, F-75015, Paris, France; Department of Clinical Haematology, AP-HP, Hôpital Necker-Enfants Malades, F-75015, Paris, France

**Keywords:** CELIAC DISEASE, GASTROINTESTINAL LYMPHOMA, GENE MUTATION

## Abstract

**Objective:** Enteropathy-associated T-cell lymphoma (EATL) is a rare but severe complication of celiac disease (CeD), often preceded by low-grade clonal intraepithelial lymphoproliferation, referred to as type II refractory CeD (RCDII). Knowledge on underlying oncogenic mechanisms remains scarce. Here, we analysed and compared the mutational landscape of RCDII and EATL in order to identify genetic drivers of CeD-associated lymphomagenesis.

**Design:** Pure populations of RCDII-cells derived from intestinal biopsies (n=9) or sorted from blood (n=2) were analysed by whole exome sequencing, comparative genomic hybridization and RNA-sequencing. Biopsies from RCDII (n=50), EATL (n=19), type I refractory CeD (n=7) and uncomplicated CeD (n=7) were analysed by targeted next-generation sequencing. Moreover, functional in vitro studies and drug testing were performed in RCDII-derived cell lines.

**Results:** 80% of RCDII and 90% of EATL displayed somatic gain-of-functions mutations in the JAK1-STAT3 pathway, including a remarkable p.G1097 hotspot mutation in the JAK1 kinase-domain in approximately 50% of cases. Other recurrent somatic events were deleterious mutations in NFκB-regulators *TNFAIP3* and *TNIP3* and potentially oncogenic mutations in *TET2, KMT2D* and *DDX3X*. JAK1 inhibitors and the proteasome inhibitor bortezomib could block survival and proliferation of malignant RCDII-cell lines.

**Conclusion:** Mutations activating the JAK1-STAT3 pathway appear to be the main drivers of CeD-associated lymphomagenesis. In concert with mutations in negative regulators of NFκB, they may favour the clonal emergence of malignant lymphocytes in the cytokine-rich coeliac intestine. The identified mutations are attractive therapeutic targets to treat RCDII and block progression towards EATL.

## INTRODUCTION

Celiac disease (CeD) is a frequent autoimmune-like enteropathy induced by gluten in genetically predisposed individuals, with a worldwide distribution and rising incidence in industrialized countries.[1] Although generally reversible and effectively controllable through gluten-free diet (GFD), CeD predisposes to severe lymphoid malignancies. These malignant complications are thought to develop from the compartment of intraepithelial lymphocytes (IEL)[2–4] and can manifest either as type II refractory CeD (RCDII), a low-grade clonal intraepithelial lymphoproliferation [5,6] or as a highly aggressive enteropathy-associated T-cell lymphoma (EATL).[7] EATL can arise *“de novo”* in CeD patients, but up to 50% of EATL develop through an intermediary RCDII step.[5,6] RCDII is characterized by the massive infiltration of the gut epithelium by lymphocytes which display clonal rearrangements of the T-cell receptor (TCR) and an unusual immunophenotype combining both T- and NK-cell traits that reflects their origin from a small subset of innate-like T-IEL.[2,8–11] Although initially localized within the gut epithelium, RCDII-IEL can disseminate into *lamina propria* and blood and ultimately reach other organs.[12] Diagnosis of RCDII is challenging and requests a combination of molecular and phenotypic methods.[5,11,13] Indeed, due to their normal cytological appearance and low proliferative rate, RCDII-IEL are difficult to differentiate from the normal polyclonal population of CD8+ T-cells that infiltrate the gut epithelium in active CeD. In contrast, EATL are characterized by a pleomorphic infiltrate of medium and large-sized proliferative lymphoma cells[14] that usually express integrin CD103, a signature of their intraepithelial origin,[4,15] and CD30, an activation marker absent in RCDII-IEL.[7,16] EATL prognosis is very poor, with a 5-year survival rate of around 60% in *de novo* EATL, which decreases to less than 5% in patients with EATL complicating RCDII,[7] stressing the need for strategies allowing early diagnosis and efficient treatment of RCDII in order to prevent its transformation into EATL. Based on the analysis of TCRγ rearrangements, we have reported that EATL complicating RCDII share a common clonal origin with RCDII-IEL.[3] Mechanisms of IEL transformation remain however largely elusive although some genomic alterations have been reported in a small number of RCDII[10] and EATL[17] cases.

Herein, we have applied a combination of genetic approaches in order to comprehensively map the mutational landscape of RCDII and EATL, to identify genetic events driving lymphocyte transformation during RCDII and its progression to EATL, and to delineate potential differences between EATL complicating RCDII and those arising *de novo* in CeD. We demonstrate the outstanding frequency of *JAK1* and *STAT3* GOF mutations but also reveal the frequent occurrence of deleterious mutations in negative regulators of NFκB and in several epigenetic regulators, thus revealing candidate targets for therapeutic intervention.

## METHODS

### Tissue sampling

Tissue samples and blood were obtained from patients with CeD (n=7), RCDI (n=7) and RCDII (n=50) enrolled prospectively in the French national “centres experts des lymphomes associés à la maladie coeliaque” (CELAC) registry until June 2018. Nineteen patients had EATL complicating CeD (n=8) or RCDII (n=11). Paired tumour and non-tumour biopsies were available for 12/19 patients (CeD=3, RCDII=9). Clinical characteristics are summarized in **Table 1** and **Supplementary Table 1**. For detailed description of diagnostic procedures see **Supplementary Experimental Procedures**.

**Table 1:**
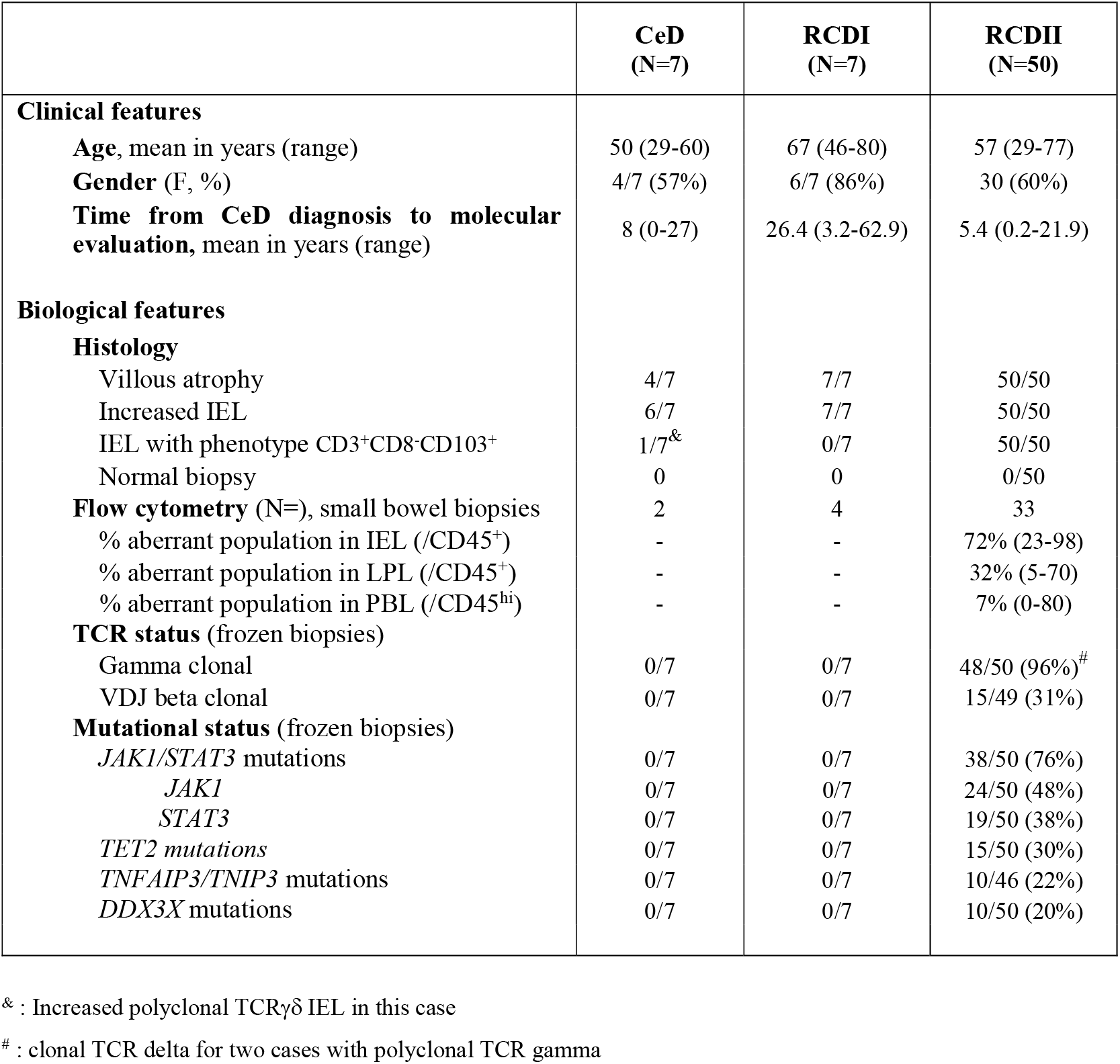
Characteristics of CeD, RCDI and RCDII patients

### Genomic analyses

Genomic DNA was extracted from cell lines, peripheral blood mononuclear cells, and frozen biopsy specimens with QIAamp DNA Mini Kit according to the manufacturer’s instructions. Comparative genomic hybridization (CGH), whole exome sequencing (WES), targeted next-generation sequencing (TNGS) and targeted amplicon sequencing (TAS), TCR rearrangements were performed and analysed as detailed in **Supplementary Experimental Procedures**.

### Cell culture and drug inhibition assays

Patient-derived cell lines were obtained as described[8] and maintained in complete RPMI (Invitrogen, Thermo Fisher Scientific, Villebon-sur-Yvette, France) with 20ng/ml IL-15 (R&D Systems, Bio-Techne, Lille, France). Cells were incubated or not with various drugs, were analyzed by flow cytometry for apoptosis and proliferation assays, by western blot for STAT3 phosphorylation determination or imaging flow cytometry for nuclear NFκB/p50 translocation. For a detailed description see **Supplementary Experimental Procedures**.

### Statistical Analysis

Prism 6 software (GraphPad Software Inc., La Jolla, CA, USA) was used for statistical analysis. Multiple comparisons were performed via One-way ANOVA with Dunnett’s correction, and categorical comparisons were analysed with Fisher’s exact test. Survival analysis was performed using Kaplan Meier Curves and Log Rank test in patients followed up from diagnosis to the latest news in June 2018.

## RESULTS

### Whole exome sequencing and comparative genomic hybridization reveal recurrent mutations in RCDII

In order to establish a comprehensive and complete catalogue of the somatic genetic events associated with malignant transformation in RCDII, WES and CGH were performed on pure populations of RCDII-cells (>99% sCD3-iCD3+sTCR-) that were either cell lines derived from biopsies of RCDII patients (n=9) or FACS-sorted circulating RCDII-cells from peripheral blood (n=1; case 3). For WES, RCDII-cells were compared to normal autologous CD3+ PBL (n=9) or sorted T-cells (n=1, case 3).

After curation of WES data, the number of non-synonymous somatic mutations ranged from 39 to 110 (median=67) per case. (**Figure 1A; Supplementary Figure 1**). Most frequent curated exonic mutations were C>T transitions (32%), followed by C>A transversions (23%), except for patient 8 (P8) which showed mainly T>G transversions (**Figure 1B**). Integrative analysis of WES and CGH disclosed both recurrent and distinctive somatic alterations in RCDII-cells. Predicted GOF somatic *JAK1* and or *STAT3* missense mutations were most frequent, occurring in 7/10 and 4/10 cases respectively. *STAT3* mutations were predominantly found within the SH2-domain and *JAK1* mutations in the JH1-kinase-domain. In one case, a *JAK1* frameshift mutation was observed on the second allele and, in 3 additional cases, CGH revealed concomitant loss-of-heterozygosity (LOH) of a region including *JAK1* on chromosome 1p (**Figure 1C&D**). Two patients carried both *JAK1* and *STAT3* mutations and three cases with only *JAK1* or *STAT3* mutation showed additional deleterious alterations in negative regulators of the JAK-STAT pathway such as CISH, SH2B3 or SOCS3[18]. Of note, the only case without *JAK1* or *STAT3* mutation displayed deleterious homozygous mutations of *SOCS1* as well as of *SH2B3*, resulting in 100% prevalence of mutations activating the JAK1-STAT3 pathway[18]. Additional recurrent somatic events included mutations of the X-linked RNA-helicase DDX3X[19] (5/10), of the methyl-cytosine dioxygenase *TET2[20]* (3/10) as well as of the tumour suppressor *PRDM1/BLIMP1*[21] (3/10).

**Figure 1:**
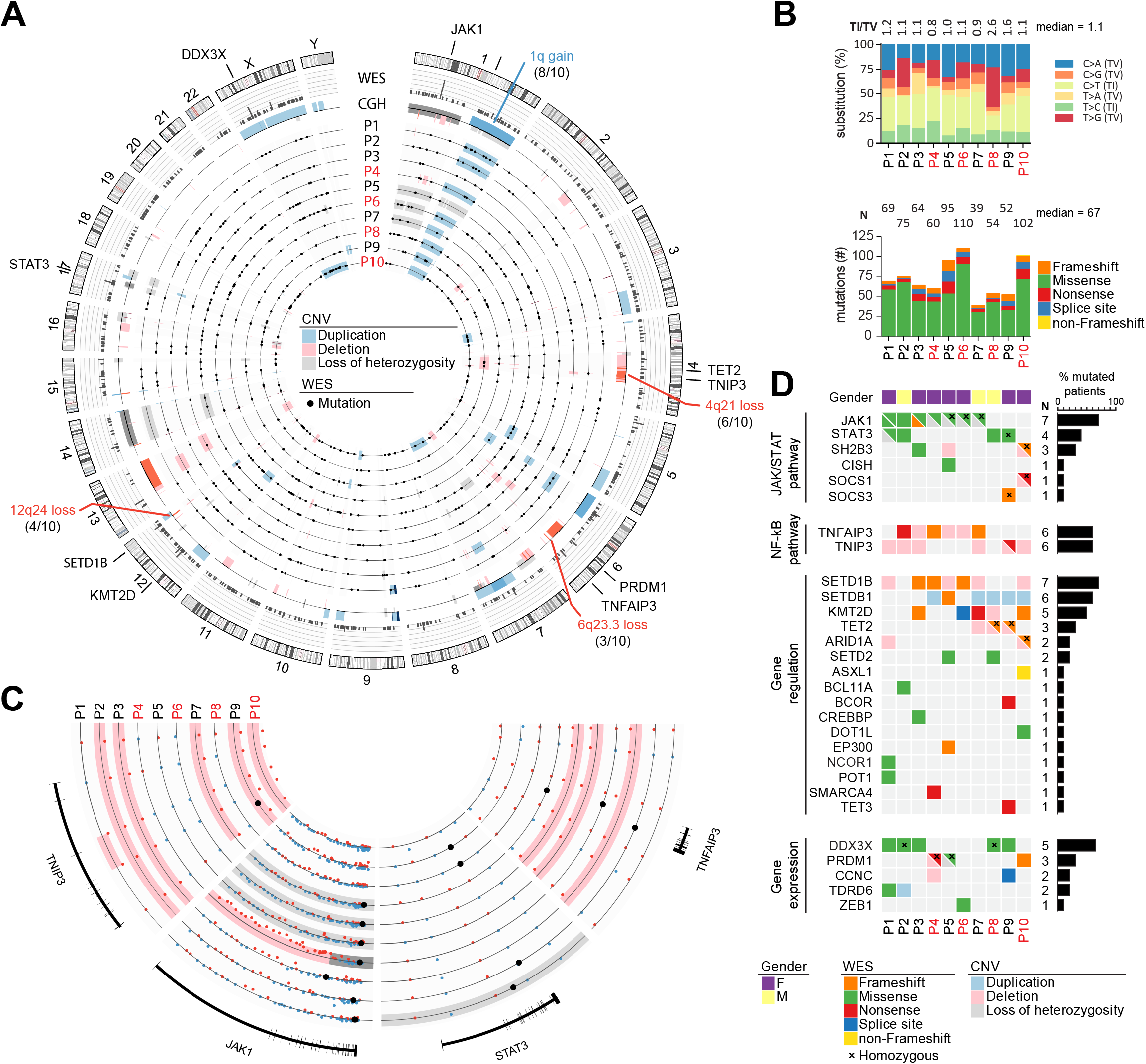
Genomic characterization of the mutational landscape of RCDII cells. Results from WES and CGH in primary RCDII-IEL lines (n=9) and RCDII cells sorted from peripheral blood (n=1). (**A**) Circos plot depicts distribution of curated somatic variants and copy number variants (CNV) across chromosomes. Outer ring shows ideograms of human chromosomes 1-22, X and Y from p- to q-region, divided by centromeres in red and with cytogenetic bands depicted by light and dark shades. Gene names and CNV adjacent to ideograms highlight selected candidate genes or regions with lines pointing to their approximate location on the genome. Bar graphs on 2nd ring and box plot on 3rd ring summarize variants per gene and CNV per location respectively. Numbered concentric circles show WES results from individual patients as black dots and CNV as boxes colour coded as indicated in legend. (**B**) Bar graphs show proportion of transitions and transversions (upper graph) and mutational categories per patient (lower graph). (**C**) Circos plot excerpt shows magnified regions from chromosomes 1, 4, 6 and 17 with indicated gene loci for individual patients. Dots indicate hybridization status of CGH probes (blue=positive, red=negative) or small mutations (black) and coloured boxes summarize CNV coded as in A. (**D**) Heatmap summarizes somatic gene variants in combination with CNV results per patient (column) and gene (row) grouped by pathway or function with mutations colour coded as indicated in the legend. Bar graphs indicate percentage of occurrence per gene.

CGH revealed recurrent trisomy 1q (8/10) and losses within 4q (6/10) and 6q (5/10) (**Figures 1A, B&C**). Chromosome 4q losses spanned 4q21, thereby including *TNIP3*, a member of the ubiquitin editing complex for NFκB regulation.[22] The 6q losses included the well-known tumour suppressor *TNFAIP3/A20*, [23] a binding partner of TNIP3, in three cases. Deleterious somatic mutations in this gene were detected in three additional cases by WES. Overall, loss-of-function (LOF) alterations of *TNIP3* and *TNFAIP3/A20* were found in 90% (9/10), making NFκB signalling the second most affected pathway in RCDII cell lines. Other frequently affected genes included histone-modifiers, with deleterious *SETD1B* alterations (7/10) due to frameshifts (3/7) and 12q losses (4/7) including a minimal 960 Kb 12q24.31 microdeletion, gain of *SETDB1* (6/10), through chromosome 1q gain in 5 cases, as well as deleterious mutations of *KMT2D* (5/10).[24,25] Beyond this core mutational signature, RCDII-cells contained a broad spectrum of additional uniquely occurring pathogenic mutations that potentially shaped individual identity. Accordingly, transcriptional analysis of the RCDII-cell lines (n=4) revealed a heterogeneous transcriptional profile (**Supplementary Figure 2**). Importantly, transcriptional analysis (n=5) did not reveal any fusion transcripts (**data not shown**).

Since cell-lines might acquire mutations during cell culture, WES was compared between one RCDII-line derived from an intestinal biopsy and circulating RCDII cells freshly sorted from autologous peripheral blood (**Case 4 Figure 1** and **Supplementary Figure 3**). Both samples displayed the same clonal TCR-rearrangement (**data not shown**), attesting their clonal origin from a common ancestor. Many mutations, notably those involving *JAK1, TNFAIP3* and *SETD1B* overlapped between the two samples, arguing against artificial induction of these mutations during cell culture. Moreover, biopsy-derived and PBL-sorted RCDII-cells also showed substantial differences, suggesting clonal evolution and selection.

Overall, these data highlight the malignant nature of RCDII and illustrate a unique mutational profile. They suggest a driver role for *JAK1-STAT3* GOF mutations and a contribution of NFκB activating mutations to RCDII pathogenesis.

### Analysis of primary intestinal biopsies confirms recurrent activating mutations of the JAK1-STAT3 and NFκB pathways

We next screened frozen biopsies from 50 RCDII patients via targeted next-generation sequencing (TNGS) of a panel of 104 genes involved in T-cell malignancies (**Supplementary File 1**). Results were confirmed using targeted amplicon resequencing (TAS) that covered mutated regions identified by TNGS and 22 additional genes identified in the explorative WES- and CGH-based approach (**Supplementary File 2**). TNGS and TAS were chosen over WES and CGH to analyse biopsies as both techniques allow deeper sequencing and were therefore anticipated to detect RCDII-associated mutations despite the low frequency of RCDII-cells in biopsies. Duodenal biopsies from uncomplicated CeD (n=7) and RCDI (n=7) patients served as controls. Description of patients and controls is provided in **Table 1**.

All biopsies from CeD and RCDI controls showed a polyclonal TCRγ- and VDJβ-repertoire. In contrast, 48/50 (96%) of RCDII biopsies displayed clonal TCRγ-rearrangement while 2/50 (4%) had polyclonal TCRγ but clonal TCRδ. Abnormal expansion of RCDII-cells was confirmed by flow cytometry in 30/33 patients. Three patients had RCDII-IEL that lacked CD4 and CD8 but showed expression of surface CD3 with either TCRγδ (2 cases with 70% and 95% TCRγ+ IEL respectively) or TCRαβ (95%).

While no mutation was detected in CeD or RCDI, a mean of 3.9 mutated genes (0-10) was observed per RCDII biopsy (**Figure 2A**). Again, the JAK-STAT pathway was the most abundantly mutated signalling pathway, with frequent mutations in *JAK1* (48%) and *STAT3* (38%) as well as in the negative JAK-STAT regulators *SOCS1* (12%) and *SOCS3* (8%). Overall 43/50 (86%) patients showed at least one somatic alteration of the *JAK1-STAT3* axis. Other frequent mutations were again observed in *TET2* (30%), *KMT2D* (22%) and *DDX3X* (20%), confirming the above findings. Mutations of the NFκB regulating genes *TNFAIP3* (13%) and *TNIP3* (9%) were found in 22% (10/46) of RCDII biopsies tested. Globally, most mutations were missense, followed by nonsense and frameshift mutations (**Figure 2B**). Notably, mutations in *TNFAIP3/A20, TNIP3, KMT2D, TET2, SOCS1* and *SOCS3* were mainly nonsense and frameshift mutations indicating their deleterious nature (**Figure 2B**). There were, however, no detectable CNV in *TNFAIP3/A20* or *TNIP3*, in contrast to RCDII-lines. Four RCDII samples (8%) did not contain any detectable mutation, either because the panel did not cover the corresponding genes or due to insufficient tissue infiltration despite detection of clonal TCRγ- or TCRδ-rearrangement within the same biopsy **(Figure 2A)**. The variant allele frequency (VAF) of mutations across RCDII samples ranged from 28% to 1% (**Supplementary Figure 4**), overall suggesting that the number of mutations might be underestimated, most notably copy number variants. VAF of distinct mutations were also highly variable within individual samples, suggesting intra-tumour heterogeneity. VAF of *JAK1* mutations were largely dominant in individual samples arguing for a founding role, whereas VAF of *STAT3* or *TNFAIP3/A20* and *TNIP3* mutations were often found close to average the VAF, suggesting secondary driver events. Accordingly, samples with *JAK1-STAT3* double-mutations often showed lower VAF for *STAT3* than for *JAK1* mutations **(Supplementary Figure 4)**. In RCDII, *JAK1* mutations tended to be mutually exclusive with *STAT3* and *TET2* mutations (**Figure 2C**, **Supplementary Table 2&3**) while *STAT3* mutations significantly co-occurred with *TET2* mutations. We also observed a striking co-occurrence of BCOR[26] and *TNFAIP3/A20* or CD58[27] mutations **(Figure 2C)**. Patients with *JAK1* mutations showed significantly higher frequencies of RCDII-cells among IEL and LPL (**Supplementary Table 2**), while patients with mutations in *TNFAIP3/A20* and *TNIP3* had significantly increased RCDII-cells in LPL and even showed malignant cell dissemination to the PBL compartment (**Supplementary Table 3**). We found no significant prognostic impact on overall survival in relation to the *JAK1 or STAT3* mutational status (**Supplementary Figure 5**). Contrary to NK/T-cell lymphomas,[19] *DDX3X* mutations were not associated with an inferior prognosis in RCDII patients.

**Figure 2:**
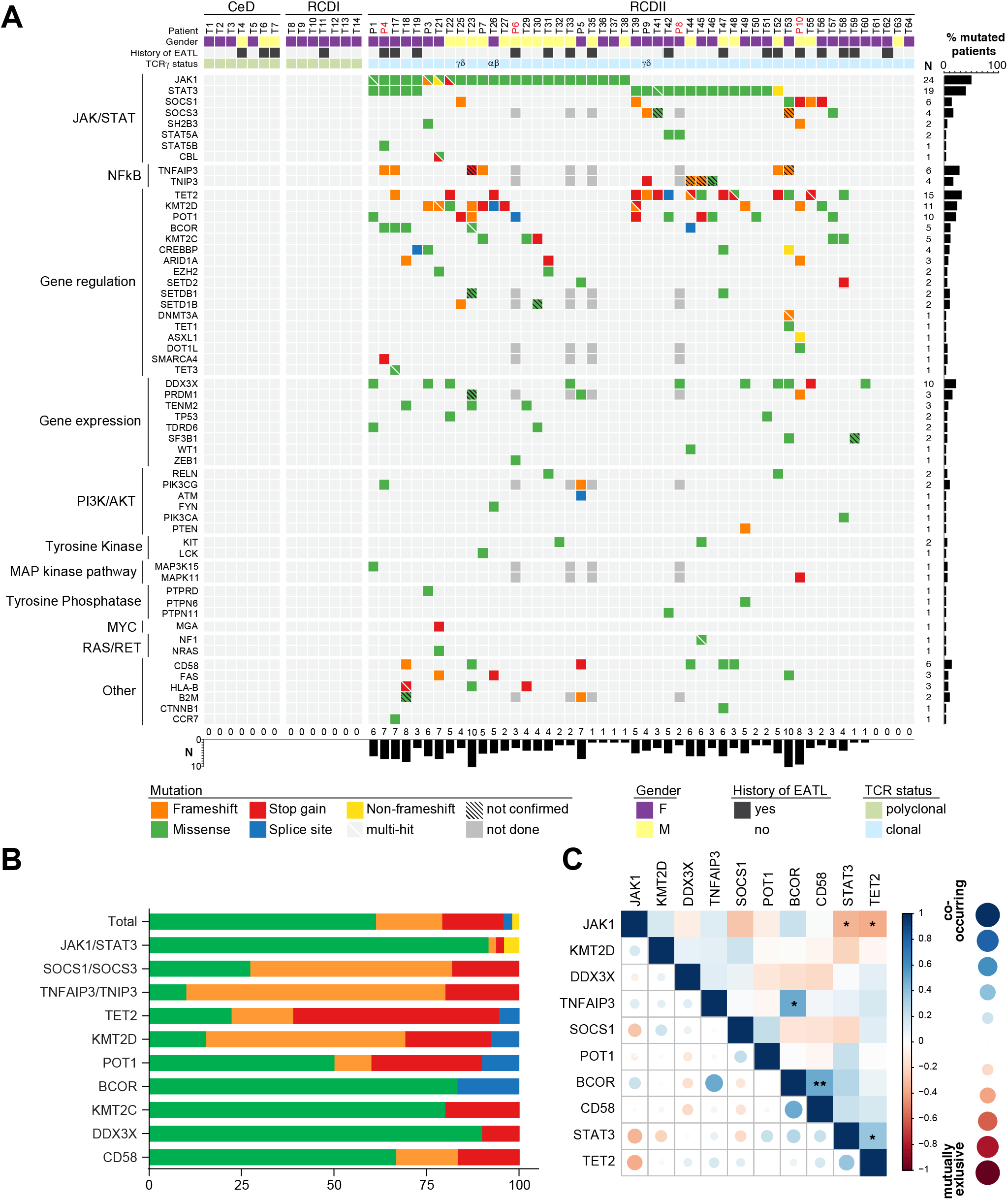
Characterization of RCDII-associated somatic mutations in intestinal biopsies. (**A**) Heatmap summarizes mutations determined by TNGS and TAS for individual patients (column) and genes (row), grouped by pathway or function. Mutations are colour coded according to the type of mutation and upper header bar shows sample ID and colour codes for gender, history of EATL and TCRγ-status as indicated in the legend. Vertical bar graph illustrates mutation frequency per gene and horizontal bar graph shows number of mutations per patient. (**B**) Stacked bar graph summarizes distribution of mutational classes according to the colour code indicated in A for selected genes. (**C**) Correlation plot visualizes co-occurrence for top 10 mutated genes in RCDII. The Pearson correlation value is coded by colour (upper right) or colour and size (lower left) as in scale on right; Significance levels were assessed via Fisher’s exact test (***=p<0.001, **=p<0.01, *=p<0.05).

Taken together, these data confirm the importance of the JAK1-STAT3 and NFκB pathways as founders or drivers of RCDII pathogenesis in a large series of primary patient samples.

### RCDII is characterized by increased cytokine responsiveness through mutations activating the JAK1-STAT3 and NFκB pathways

Topographic mapping of the recurrent *JAK1* and *STAT3* mutations indicated predominance in the SH2-domain of STAT3 (65%) and showed that virtually all (93% 26/28) *JAK1* mutations clustered at the p.G1097 position in the C-terminal JH1-kinase-domain, a highly conserved position, and the site of interaction of the JAK1 negative regulator, SOCS1[28] (**Figure 3A&B**). Within this hotspot, the p.G1097D (14/24) variant was most frequent, followed by p.G1097V/C/S/R (25%, 8%, 4% and 4% respectively). Of note, almost all JAK1 or STAT3 mutations not located in the JAK1 p.G1097 hotspot or in the STAT3 SH2-domain co-occurred with a JAK1 p.G1097 hotspot mutation. Exceptions included one case with the known STAT3 p.Q344H GOF mutation in the DNA-binding domain, and one with two STAT3 p.D173E/p.F174I mutations on the same allele. Mutations in TNFAIP3/A20 and TNIP3 were scattered over their N-terminal or central regions (**Figure 3C**). Western-blot analyses performed in RCDII-lines revealed constitutive STAT3 phosphorylation in P4 cell line with JAK1 p.G1097D mutation and loss of heterozygosity, suggesting intrinsic activation of the JAK1-STAT3 pathway. Cells from P6 with homozygous JAK1 p.G1097D and P8 with heterozygous STAT3 p.D661Y mutation as well as cells from P4 displayed enhanced and sustained phosphorylation in response to extrinsic IL-15 stimulation when compared to normal T-cell lines. This held also true for P10-derived cells, which harboured deleterious *SOCS1* and *SH2B3* mutations, although to a lesser extent (**Figure 3 D**). Analysis of NFκB activation by imaging flow cytometry revealed constitutive nuclear translocation of the NFκB/p50 subunit in the RCDII-lines derived from P5 and P6, with loss of one *TNFAIP3/A20* allele (**Figure 3E & Supplementary Figure 6**) and enhanced TNFα-induced nuclear translocation in the RCDII-lines from P4, with a c.102_103insGG/p.G35 frameshift in *TNFAIP3/A20*, and from P10, with loss of one *TNIP3* allele, when compared to the P8 cell line, with wild-type alleles for these genes (**Supplementary Figure 6**).

**Figure 3:**
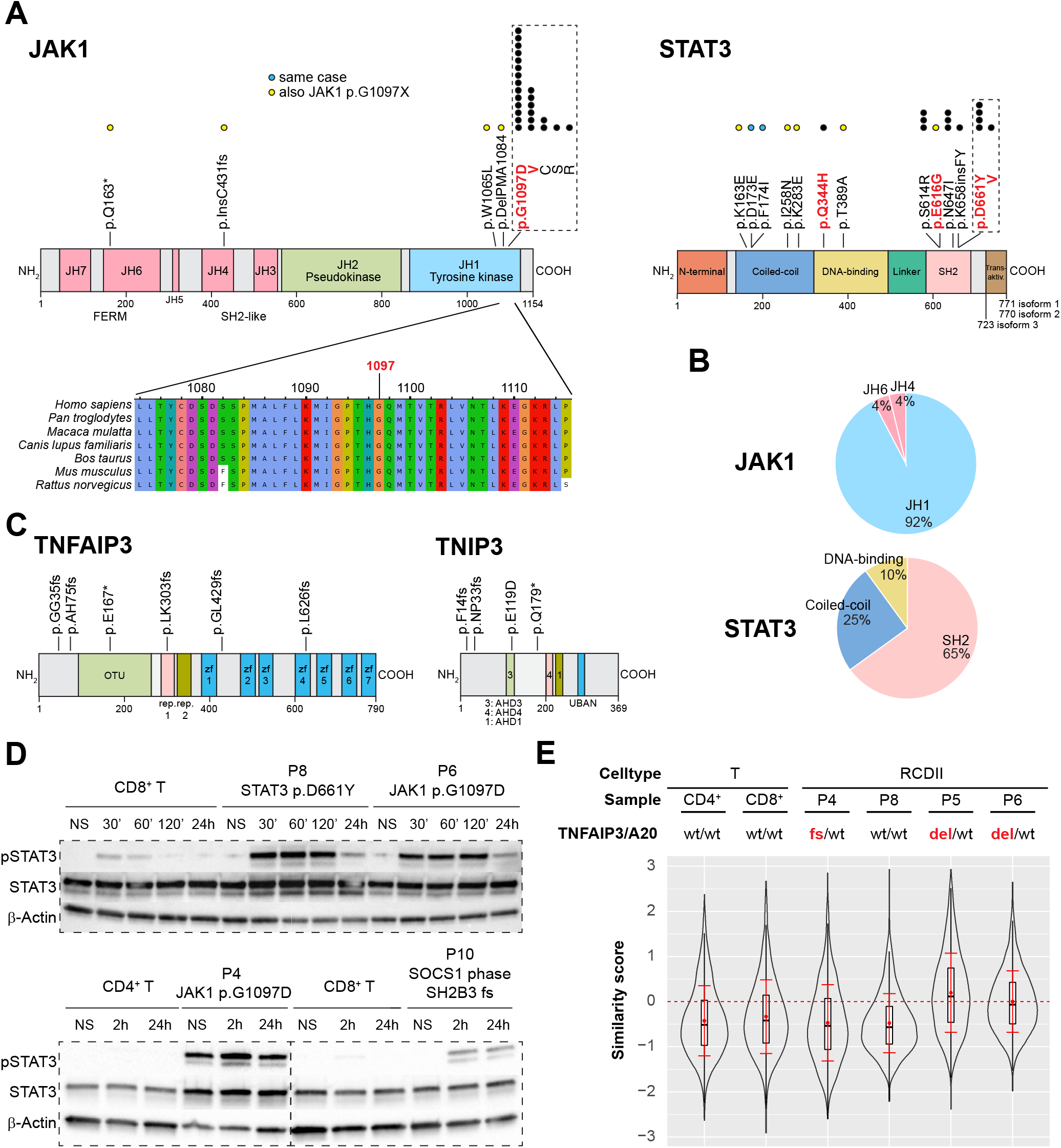
Functional analysis of JAK1-STAT3 and NFκB pathway activating mutations in RCDII cell lines. (**A**) Schematic illustration of topographic localization of JAK1-STAT3 mutations found in RCDII and conservation plot for indicated species centred on the JAK1 p.G1097 hotspot. Dots represent mutations per position with yellow dots depicting those co-occurring with JAK1 p.G1097 hotspot mutations and blue dots for mutations found in the same patient. Mutations highlighted in red indicate known GOF variants. (**B**) Pie charts show relative distribution of mutations per protein domain for JAK1 and STAT3. (**C**) Topographic localization of TNFAIP3/A20 and TNIP3 mutations found in RCDII (**D**) Western blots for pSTAT3, STAT3 and β-actin for RCDII-cell lines from 4 patients and control CD3+ T-cell lines upon stimulation with 20ng/ml IL-15 for indicated time points; NS=not stimulated. (**E**) Violin plot shows translocation scores for NFκB/p50 in unstimulated RCDII-cell lines (n=4) and in unstimulated control CD3+CD4+ and CD3+CD8+ T-cell lines. Representative results from at least two independent experiments.

Altogether, these data indicate that RCDII cells contain highly recurrent mutations in the JAK1-STAT3 and NFκB pathways that result in their intrinsic activation and/or enhanced responsiveness to extrinsic inflammatory stimuli.

### The oncogenic signatures of EATL and RCDII largely overlap and are dominated by JAK1-STAT3 mutations

EATL can complicate RCDII and, thereby, share the same clonal origin as RCDII IEL[3] (RCDII-EATL) but it can also develop in CeD patients without RCDII (“de novo” EATL). Prognosis of EATL is better in the latter case,[7] raising the possibility that distinctive oncogenic events underlie these two entities. To address this hypothesis, we compared TNGS results in biopsies obtained from RCDII-EATL (n=11) and *de novo* EATL (n=8).

Overall, the mutational profile of both RCDII-EATL and *de novo* EATL largely overlapped with that of RCDII, including recurrent GOF mutations at the JAK1 p.G1097 hotspot and in the SH2-domain of STAT3, as well as *SOCS1* frameshift or nonsense mutations in two patients without *JAK1* or *STAT3* mutations (**Figure 4**). 37% of EATL (9/19) showed multiple *JAK1* or *STAT3* mutations with up to three mutations in two patients and *JAK1* and *STAT3* double-mutations in 6/19 (32%) patients. Other mutations frequently found in EATL were in *KMT2D* (37%), *TET2* (32%), *DDX3X* (32%), *TNFAIP3* (28%) and POT1 [29] (26%). Interestingly, the mean number of mutations detectable by TNGS and TAS were comparable between *de novo* EATL and RCDII-EATL (**Supplementary Figure 7**). There were, however, some differences observed as all *de novo* EATL but only 55% of RCDII-EATL showed at least one p.G1097 *JAK1* mutation (p=0.045), whereas *STAT3* mutations tended to be overrepresented in RCDII-EATL (63.6%) when compared to *de novo* EATL (25%; p=0.17). Moreover, mutations in *TNFAIP3* (p=0.036), *BCOR* and *SMARCA4* were only found in RCDII-EATL, while *PRDM1* mutations were exclusively detected in *de novo* EATL (3/8).

**Figure 4:**
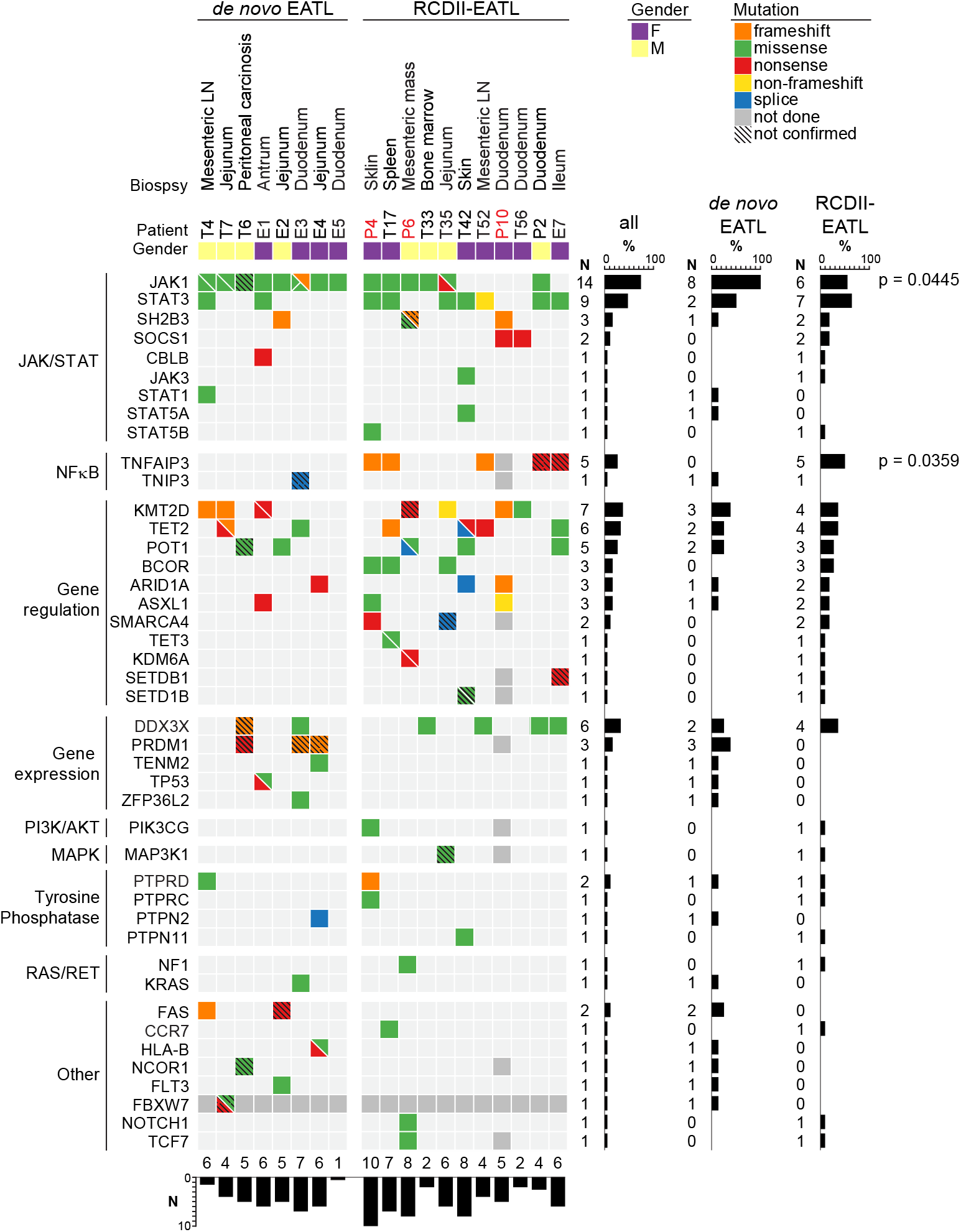
Mutational profiles of EATL complicating RCDII or developing *de novo* in celiac disease. Heatmap summarizes mutations determined by TNGS and TAS for individual patients (column) with *de novo* EATL (left block) or RCDII-EATL (right block). Genes (row) are grouped by pathway or function. Mutations are colour coded according to the type and upper header bar shows sample ID and colour codes for gender as indicated in the legend below. Horizontal bar graph illustrates frequency of mutations per gene with adjacent absolute numbers for all EATL, *de novo* EATL and RCDII-EATL. Vertical bars illustrate absolute counts of mutations per patient. p-values are shown for categorical differences between *de novo* EATL and RCDII-EATL as assessed via Fisher’s exact test.

We also compared RCDII-EATL (n=9) to autologous RCDII biopsies without evidence of EATL (**Figure 3A&4**). Most RCDII-EATL samples showed increased VAF when compared to their non-EATL controls (**Figure 5**). Strikingly, *JAK1-STAT3* and *TNFAIP3/A20* mutations were never lost during progression from RCDII to EATL, supporting their founding or driving role in transformation. Of note, EATL in P4 showed particular expansion of a STAT3 p.E616G mutated clone (**Figure 5**), a variant that was not detected in the autologous RCDII-cell line, pointing to the co-existence or emergence of various clones. When compared to autologous RCDII biopsies without EATL, RCDII-EATL samples contained additional pathogenic mutations in *STAT3, SH2B3, JAK3, KMT2D, BCOR, ARID1A, SETD1B, PTPRC, PTPRD, NF1* and *NOTCH1*, suggesting that these newly acquired mutations represent driver events supporting RCDII transformation into EATL. Moreover, *JAK1-STAT3* double-mutations as well as *TNFAIP3/A20* mutations tended to predispose to EATL transformation (**Supplementary Figure 8**). Definitive demonstration of the prognostic value of these mutations to predict the risk of RCDII progression to EATL will however require analysis of more cases.

**Figure 5:**
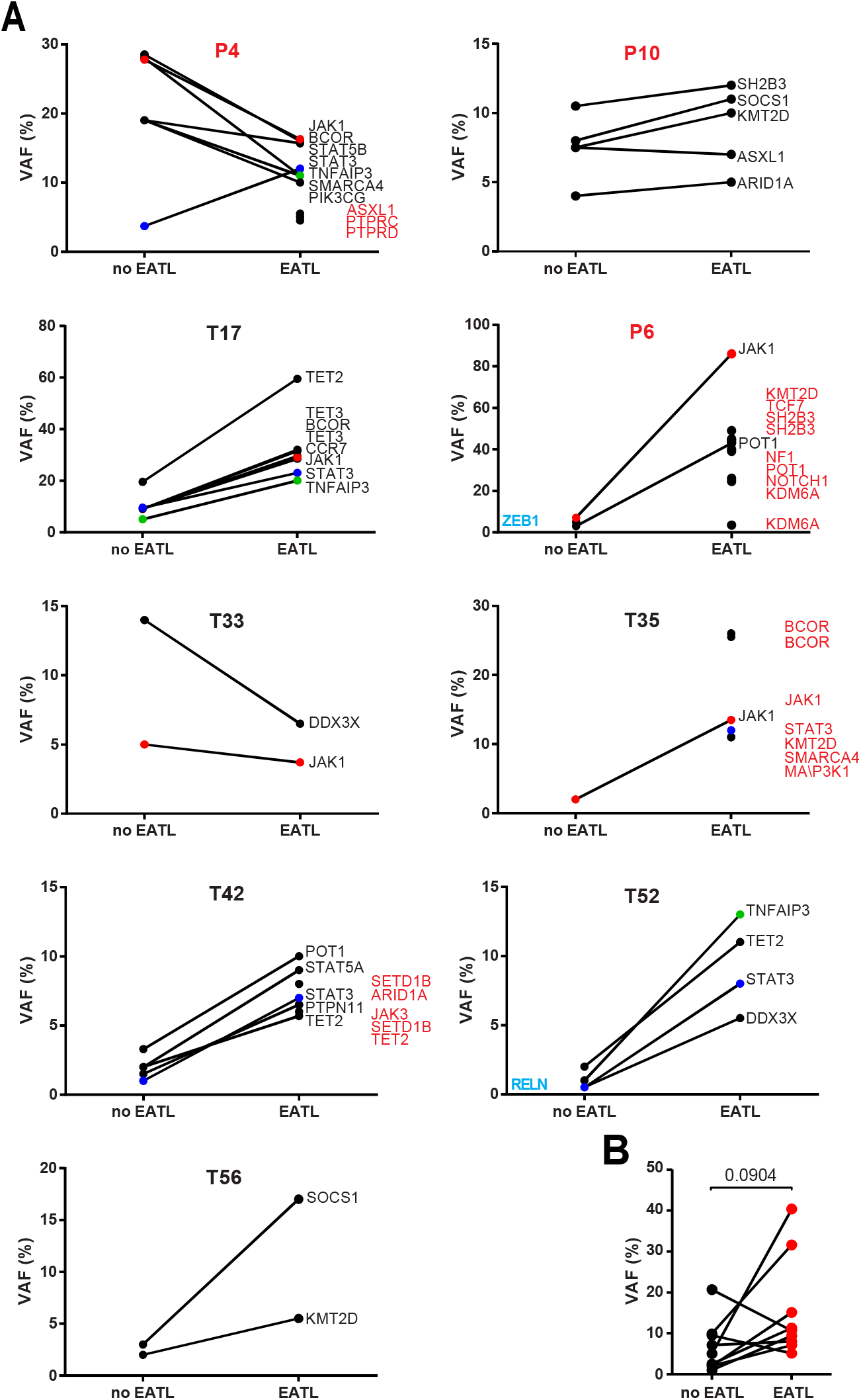
Comparison of somatic mutations during transformation of RCDII into EATL. (**A**) Before and after plots show mean VAF of individual mutations of RCDII samples without (no EATL) or with EATL (EATL) for individual patients. Highlighted genes were detected in only one group (blue=no EATL, red=EATL). (**B**) Before and after plot summarizes mean variant allelic frequencies (VAF) of RCDII samples without (no EATL) or with EATL (EATL) in each patient; p-value was calculated via paired two-tailed t-test.

Overall, these data indicate that common mechanisms underlie lymphomagenesis in *de novo* EATL and RCDII-EATL, but with restriction of *TNFAIP3/A20* mutations to RCDII-EATL, whereas *JAK1* mutations were overrepresented in *de novo* EATL.

### The JAK1-STAT3 pathway is a potential therapeutic target in RCDII

Given the severe prognosis of EATL, identifying therapies that efficiently treat RCDII and block progression to EATL is indispensable. RCDII-lines were used as *in vitro* preclinical models to assess the therapeutic efficacy of drugs targeting the highly recurrent *JAK1-STAT3* GOF mutations using ruxolitinib, which inhibits JAK1 and JAK2, and abrocitinib (PF-04965842), a specific JAK1-inhibitor.[30] Their effect was compared to that of budesonide, a corticosteroid commonly used in RCDII[31] and bortezomib, a proteasome inhibitor which has demonstrated efficacy in myeloma by modulating survival and apoptosis of malignant cells[32] and which has also been shown to interfere with STAT3 signalling.[33–35] Confirming and extending our published results in two distinct RCDII-lines, [10] both ruxolitinib and abrocitinib reduced proliferation, induced apoptosis and, simultaneously, inhibited STAT3 phosphorylation in all 4 RCDII-cell lines tested (**Figure 6A&B**). These drugs, however, exerted comparable or even stronger effects in cultured control T-cells. Similarly, budesonide impaired survival and growth of both RCDII- and normal T-cell lines. Moreover, and in line with the lack of clinical response of P6 to budesonide, this drug had no effect on the RDCII-line derived from intestinal biopsies of P6. In contrast, the reversible 26S proteasome inhibitor bortezomib exerted pro-apoptotic and/or anti-proliferative effects on the 4 RCDII-lines, while normal T-cell lines remained largely unaffected. Moreover, bortezomib was also able to inhibit STAT3 phosphorylation in the 4 RCDII-lines **(Figure 6B**).

**Figure 6:**
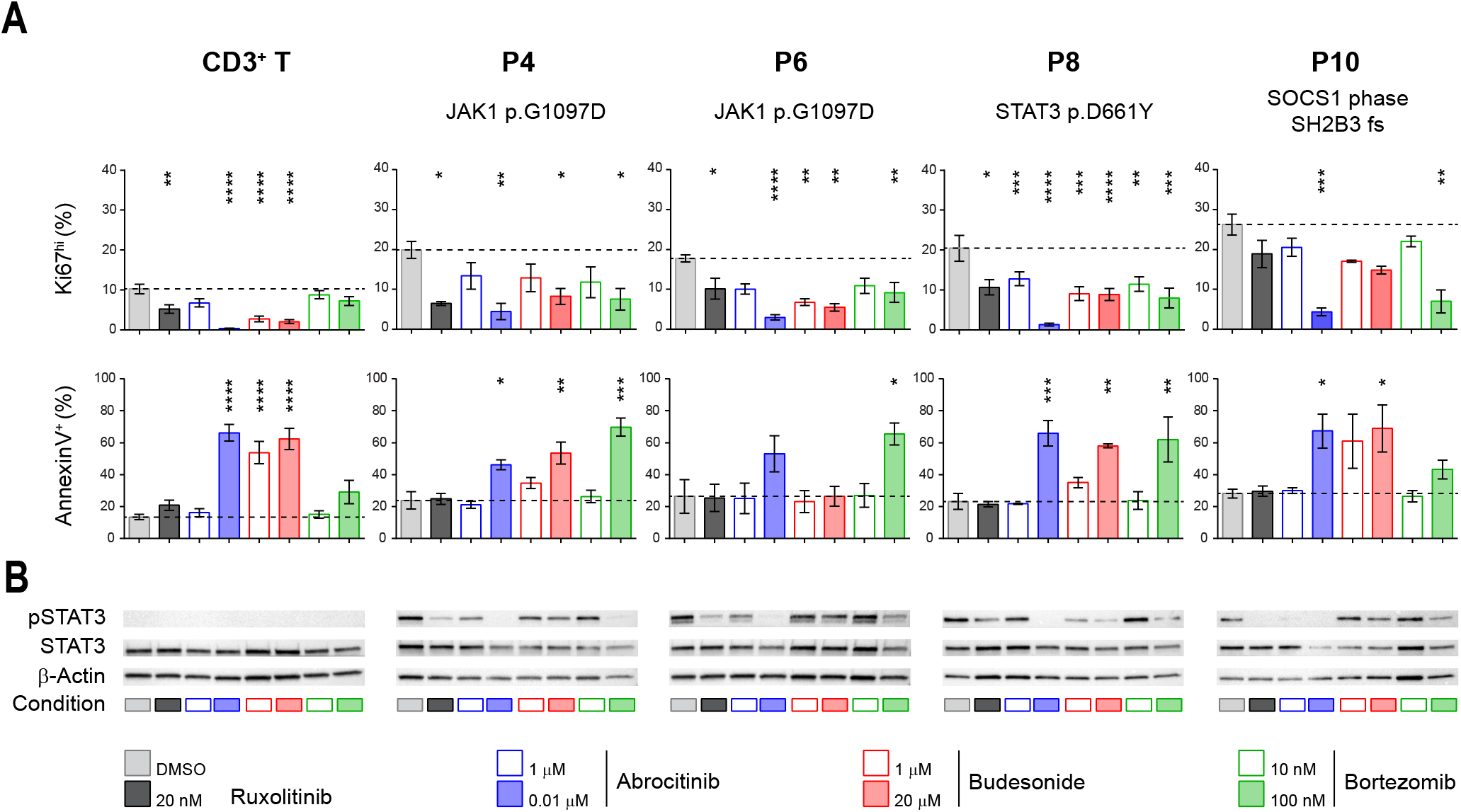
Differential efficacy of candidate therapeutic drugs in RCDII cell lines. (**A**) Bar plots show mean percentages +/− standard deviation of flow cytometry based assessment of AnnexinV+ (left column) and Ki67^hi^ cells (right column) from four patients (n=3-5) and control CD3+ T-cells (n=14) after 72h of the indicated treatment; asterisks denote statistical significant change relative to untreated (DMSO) condition; p-values (****=p<0.00001, ***=p<0.001, **=p<0.01, *=p<0.05). (**B**) Representative Western blots for pSTAT3, STAT3 and β-actin for RCDII-cell lines from four patients and T-cells as controls upon 24h of treatment with indicated drugs or vehicle (DMSO).

Overall these functional studies emphasize the relevance of JAK1-inhibitors for the treatment of RCDII and EATL. They also reveal the capacity of the proteasome inhibitor bortezomib to eliminate malignant cells while concomitantly preserving normal T-cells.

## DISCUSSION

This study, based on the largest cohort of RCDII and EATL patients to date, identifies *JAK1* and *STAT3* mutations in the vast majority of the 50 RCDII and 19 EATL studied, strongly supporting their driver role in CeD-associated lymphomagenesis. Almost all detected *JAK1* and *STAT3* variants have been reported as GOF mutations[10,36,37] and, completing our previous report,[10] additional RCDII-cell lines showed constitutive and/or enhanced cytokine-driven phosphorylation of STAT3. Frequent deleterious mutations of the negative JAK-STAT regulators and, notably, of SOCS1 in the rare cases without *JAK1* or *STAT3* mutations, further stress the outstanding role of this pathway in CeD-associated oncogenesis. A role of the JAK-STAT pathway has already been highlighted in other intestinal lymphomas. Thus, recurrent *JAK3* and *STAT5* GOF mutations have been observed in monomorphic epitheliotropic intestinal T cell lymphoma (MEITL), a highly aggressive lymphoma that is not associated with CeD[17,38] and *JAK2-STAT3* fusion transcripts associated with STAT5 activation have been reported in indolent intestinal CD4+ T-cell lymphomas.[39] We now show that *JAK1* and *STAT3 GOF* mutations are a hallmark of CeD-associated lymphomas. Moreover, the JAK1 p.G1097 hotspot identified in almost 50% of the cohort has been rarely reported in any other malignancies to this extend[36] and is therefore a potential diagnostic marker for CeD-associated lymphomas. This is in line with data reported in a small number of EATL[17] Interestingly, mutations in negative regulators of NFκB were detected in almost all RCDII-cell lines or RCDII-cells sorted from blood. They were detected less frequently in biopsies of RCDII and RCDII-EATL. Many alterations were copy number variations and may not have been detected by NGS in biopsies. The tumour-suppressive function of TNFAIP3/A20 is well described.[23] The role of TNIP3, a partner of TNFAIP3/A20 in the ubiquitin editing complex[22] is much less known, but TNIP3 inversion was reported in one case of indolent intestinal CD4+ T lymphoma. [40] CeD pathogenesis has been mainly linked to cytokines such as IFNγ, IL-15 or IL-21 that trigger the JAK-STAT pathway.[41,42] Yet, some studies have pointed to a potential role of NFκB signalling. Predisposing *TNFAIP3/A20* and *REL* gene risk variants have been associated with CeD[43]. TNFα, a potent activator of NFκB, was shown to be produced by IEL[44] and gliadin-specifìc CD4+ T-cells and could promote the growth of RCDII-lines *in vitro.* [45] Mutations activating the JAK1-STAT3 and NFκB pathways may thus synergize and, in conjunction with cytokines released in the inflammatory CeD intestine, promote the clonal expansion of malignant RCDII-cells as well as stimulate their autonomous production of cytokines and their cytotoxicity against epithelial cells,[8,9] overall creating the vicious circle of a genotoxic inflammatory environment that favours genomic instability, enabling accumulation of more genetic aberrations and ultimately leading to transformation into EATL. Accordingly, other potentially oncogenic mutations were observed in epigenetic modifiers such as TET2[20], KMT2D[24] and in the translational regulator DDX3X.[19]

The overlap between the mutational fingerprints of individual RCDII cases and their corresponding EATL strongly supports the sequential model for EATL development through an intermediate RCDII phase. The comparable mutational profile of *de novo* EATL and RCDII-EATL point to similar mechanisms of lymphomagenesis. One intriguing difference was the increased prevalence of J*AK1* mutations in *de novo* EATL, while RCDII-EATL showed more JAK1-STAT3 double-mutations and TNFAIP3/A20 LOF variants, which also tended to predispose to progression of RCDII to EATL. The latter result is in keeping with the previous demonstration of the cooperative effect of *STAT3* and *JAK1* GOF mutations in malignant transformation.[36] Together with the NFκB activating mutations, they may further potentiate transformation. Indeed, NFκB and STAT3 are suspected to act cooperatively to promote transcription of their target genes. [46] Moreover, NFκB signalling has been shown to be limited by SOCS1.[47]

Our results indicate that blockade of the JAK1-STAT3 pathway is one therapeutic option to inhibit the growth of malignant RCDII-cells and prevent progression into EATL. Benefits and risks however require careful consideration. JAK1-inhibitors had a profound impact on RCDII-cells but also on normal T-cell lines, which is in line with their known negative effect on the activation of cytotoxic lymphocytes, some of which may harbour anti-tumour function.[48] Conversely, JAK-inhibitors exert anti-inflammatory effects and may thereby help switch off the vicious circle that promotes CeD-associated lymphomagenesis, a benefit which could outweigh the risk of impaired tumour surveillance.[49] Interestingly, bortezomib, a drug approved for the treatment of multiple myeloma,[32] selectively impaired the growth of RCDII-cell lines. As described in myeloma, inhibition of RCDII-cells may result from the stabilizing effect of bortezomib on proteins that stimulate apoptosis and inhibit cell-cycle progression[50] but also on its inhibitory effect on STAT3 phosphorylation in RCDII-cells.[33–35] Therapies combining JAK1-inhibitors and bortezomib were shown to increase therapeutic efficacy in myeloproliferative neoplasia[51] and may thus be worthy of consideration in RCDII. Mutations identified in epigenetic modifiers provide additional clues to design personalized treatments adapted to the mutational profile of individual RCDII cases and to minimize the risk of progression to EATL.

In conclusion, the mutational landscape of CeD-associated lymphoid malignancies points to convergent mechanisms driving lymphomagenesis and EATL development. Our results support a scenario in which the cytokines present in the chronically inflamed CeD intestine contribute to the clonal outgrowth of innate-like IEL carrying highly recurrent *JAK1-STAT3* GOF mutations that synergize with mutations impairing NFκB regulation to foster transformation. Besides shedding new insight into the pathogenesis of CeD-associated lymphomagenesis, our work provides the rationale for new therapeutic strategies that may prevent RCDII progression to EATL and improve the prognosis of these most severe complications of CeD.

## Acknowledgements

The authors would like to thank all clinicians and pathologists for contributing in diagnostic approach and for help with sample management in the CELAC network:: Yoram Bouhnik, Charles-André Cuenod, Sabine Brechignac, Matthieu Allez, Jacques Cosnes, Agnès Fourmestraux, Jean-Charles Delchier, Jehan Dupuis, Corinne Haioun, Taoufik El Gnaoui, Eric Lerebours, Guillaume Savoye, Herve Tilly, Bernard Flourie, Bertrand Coiffier, Xavier Hebuterne, Nadia Arab, Jérôme Filippi, Stéphane Schneider, Frank Zerbib, Noel Milpied, Krimo Bouabdallah, Reza Tabrizi, Stéphane Vigouroux, Arnaud Pigneux, Thibaut Leguay, Marie-Sarah Dilhuydy, Charles Dauriac, Serge Bologna, Cyrille Hulin, Caroline Bonmati, Fréderic Magnin, Dana Ranta, Tamara Matysiakbudnik, Eric Deconinck, Philippe Pouderoux, Bruno Bonaz, Remy Gressin, Franck Carbonnel, Jean-Marc Gornet, Julien Branche, Georgette Saint-Georges, Jean-Marie Reimund, Stéphane Nancey, Maria Nachury, Stéphanie Viennot, Camille Zallot, Bettina Fabiani, Lysiane Marthey, Karine Juvin, Yann Le Baleur, Sandy Kwiatek, Eric Saillard, Dominique Louvel, Xavier Roblin, Philippe Beau, Pierre Feugier, Laurent Peyrin-Biroulet, Hélène Zanaldi, Hedia Brixi-Benmansour, Guillaume Cadiot, Thierry Lecomte, Jean-Francois Bretagne, Olivier Casasnovas, Denis Caillot, Laurent Bedenne, Jacques-Olivier Bay, Corinne Bouteloup, Bernard Duclos, Carine Foucaud.

## Author Contributions

Conception and design of the study: SC, VA, NCB.

Patient care and clinical data acquisition: GM, ShK, MC, DS, OH, CC.

Biological data generation and analysis: SC, LL, SB, AT, NG, PV, SK, BM, MD, MB, BT, NC, C.B-F, JB, TM, EM.

Data interpretation: SC, LL, GM, AT, VA, NCB. Drafting of the manuscript: SC, LL, AT, NCB.

All authors reviewed and approved the final manuscript.

## Patient consent

Not required

## Ethics approval

The study was approved by the Ile-de-France II ethical committee (Paris, France) with Inserm as study promoter (C08-34).

## REFERENCES

1 Ludvigsson JF, Murray JA. Epidemiology of Celiac Disease. Gastroenterol Clin North Am 2019;48:1–18. doi:10.1016/j.gtc.2018.09.004

2 Cellier C, Patey N, Mauvieux L, et al. Abnormal intestinal intraepithelial lymphocytes in refractory sprue. Gastroenterology 1998;114:471–81. doi:10.1016/S0016-5085(98)70530-X

3 Cellier C, Delabesse E, Helmer C, et al. Refractory sprue, coeliac disease, and enteropathy-associated T-cell lymphoma. Lancet 2000;356:203–8. doi:10.1016/S0140-6736(00)02481-8

4 Spencer J, Cerf-Bensussan N, Jarry A, et al. Enteropathy-associated T cell lymphoma (malignant histiocytosis of the intestine) is recognized by a monoclonal antibody (HML-1) that defines a membrane molecule on human mucosal lymphocytes. Am J Pathol 1988;132:1–5.

5 Malamut G, Afchain P, Verkarre V, et al. Presentation and Long-Term Follow-up of Refractory Celiac Disease: Comparison of Type I With Type II. Gastroenterology 2009; 136:81–90. doi:10.1053/j.gastro.2008.09.069

6 Rubio-Tapia A, Kelly DG, Lahr BD, et al. Clinical Staging and Survival in Refractory Celiac Disease: A Single Center Experience. Gastroenterology 2009;136:99–107. doi: 10.1053/j.gastro.2008.10.013

7 Malamut G, Chandesris O, Verkarre V, et al. Enteropathy associated T cell lymphoma in celiac disease: A large retrospective study. Dig Liver Dis 2013;45:377–84. doi:10.1016/j.dld.2012.12.001

8 Mention J-J, Ben Ahmed M, Bègue B, et al. Interleukin 15: a key to disrupted intraepithelial lymphocyte homeostasis and lymphomagenesis in celiac disease. Gastroenterology 2003;125:730–45. doi:10.1016/S0016-5085(03)01047-3

9 Hüe S’ Mention J-J, Monteiro RC, et al. A Direct Role for NKG2D/MICA Interaction in Villous Atrophy during Celiac Disease. Immunity 2004;21:367–77. doi:10.1016/j.immuni.2004.06.018

10 Ettersperger J, Montcuquet N, Malamut G, et al. Interleukin-15-Dependent T-Cell-like Innate Intraepithelial Lymphocytes Develop in the Intestine and Transform into Lymphomas in Celiac Disease. Immunity 2016;45:610–25. doi:10.1016/j.immuni.2016.07.018

11 Cheminant M, Bruneau J, Malamut G, et al. NKp46 is a diagnostic biomarker and may be a therapeutic target in gastrointestinal T-cell lymphoproliferative diseases: A CELAC study. Gut 2018;68:1396–405. doi:10.1136/gutjnl-2018-317371

12 Verkarre V. Refractory coeliac sprue is a diffuse gastrointestinal disease. Gut 2003;52:205–11. doi:10.1136/gut.52.2.205

13 Malamut G, Meresse B, Cellier C, et al. Refractory celiac disease: from bench to bedside. Semin Immunopathol 2012;34:601–13. doi:10.1007/s00281-012-0322-z

14 Swerdlow SH, Campo E, Harris NL, et al. Summary for Policymakers. In: Intergovernmental Panel on Climate Change, ed. Climate Change 2013 – The Physical Science Basis. Cambridge: : Cambridge University Press 2017. 1–30. doi:10.1017/CBO9781107415324.004

15 Cerf-Bensussan N, Jarry A, Brousse N, et al. A monoclonal antibody (HML-1) defining a novel membrane molecule present on human intestinal lymphocytes. Eur J Immunol 1987;17:1279–85. doi:10.1002/eji.1830170910

16 Murray A, Cuevas EC, Jones DB, et al. Study of the immunohistochemistry and T cell clonality of enteropathy-associated T cell lymphoma. Am J Pathol 1995;146:509–19.

17 Roberti A, Dobay MP, Bisig B, et al. Type II enteropathy-associated T-cell lymphoma features a unique genomic profile with highly recurrent SETD2 alterations. Nat Commun 2016;7:12602. doi:10.1038/ncomms12602

18 Seif F, Khoshmirsafa M, Aazami H, et al. The role of JAK-STAT signaling pathway and its regulators in the fate of T helper cells. Cell Commun Signal 2017;15:23. doi:10.1186/s12964-017-0177-y

19 Jiang L, Gu Z-H, Yan Z-X, et al. Exome sequencing identifies somatic mutations of DDX3X in natural killer/T-cell lymphoma. Nat Genet 2015;47:1061–6. doi:10.1038/ng.3358

20 Lio C-WJ, Yuita H, Rao A. Dysregulation of the TET family of epigenetic regulators in lymphoid and myeloid malignancies. Blood 2019;134:1487–97. doi:10.1182/blood.2019791475

21 Pasqualucci L, Compagno M, Houldsworth J, et al. Inactivation of the PRDM1/BLIMP1 gene in diffuse large B cell lymphoma. J Exp Med 2006;203:311–7. doi:10.1084/jem.20052204

22 Verstrepen L, Carpentier I, Verhelst K, et al. ABINs: A20 binding inhibitors of NF-κB and apoptosis signaling. Biochem Pharmacol 2009;78:105–14. doi:10.1016/j.bcp.2009.02.009

23 Hymowitz SG, Wertz IE. A20: From ubiquitin editing to tumour suppression. Nat Rev Cancer 2010;10:332–41. doi:10.1038/nrc2775

24 Michalak EM, Visvader JE. Dysregulation of histone methyltransferases in breast cancer – Opportunities for new targeted therapies? Mol Oncol 2016;10:1497–515. doi:10.1016/j.molonc.2016.09.003

25 Batham, Lim, Rao. SETDB-1: A Potential Epigenetic Regulator in Breast Cancer Metastasis. Cancers (Basel) 2019;11:1143. doi:10.3390/cancers11081143

26 Astolfi A, Fiore M, Melchionda F, et al. BCOR involvement in cancer. Epigenomics 2019; 11:835–55. doi:10.2217/epi-2018-0195

27 Kataoka K, Nagata Y, Kitanaka A, et al. Integrated molecular analysis of adult T cell leukemia/lymphoma. Nat Genet 2015;47:1304–15. doi:10.1038/ng.3415

28 Liau NPD, Laktyushin A, Lucet IS, et al. The molecular basis of JAK/STAT inhibition by SOCS1. Nat Commun 2018;9:1558. doi:10.1038/s41467-018-04013-1

29 Calvete O, Garcia-Pavia P, Domínguez F, et al. The wide spectrum of POT1 gene variants correlates with multiple cancer types. Eur J Hum Genet 2017;25:1278–81. doi:10.1038/ejhg.2017.134

30 Gooderham MJ, Forman SB, Bissonnette R, et al. Efficacy and Safety of Oral Janus Kinase 1 Inhibitor Abrocitinib for Patients With Atopic Dermatitis. JAMA Dermatology 2019;155:1371. doi:10.1001/jamadermatol.2019.2855

31 Mukewar SS, Sharma A, Rubio-Tapia A, et al. Open-Capsule Budesonide for Refractory Celiac Disease. Am J Gastroenterol 2017;112:959–67. doi:10.1038/ajg.2017.71

32 Mujtaba T, Dou QP. Advances in the understanding of mechanisms and therapeutic use of bortezomib. Discov Med 2011;12:471–80.

33 Baran-Marszak F, Boukhiar M, Harel S, et al. Constitutive and B-cell receptor-induced activation of STAT3 are important signaling pathways targeted by bortezomib in leukemic mantle cell lymphoma. Haematologica 2010;95:1865–72. doi: 10.3324/haematol.2009.019745

34 Lam LT, Wright G, Davis RE, et al. Cooperative signaling through the signal transducer and activator of transcription 3 and nuclear factor-κB pathways in subtypes of diffuse large B-cell lymphoma. Blood 2008;111:3701–13. doi:10.1182/blood-2007-09-111948

35 Bao X, Ren T, Huang Y, et al. Bortezomib induces apoptosis and suppresses cell growth and metastasis by inactivation of Stat3 signaling in chondrosarcoma. Int J Oncol 2017;50:477–86. doi:10.3892/ijo.2016.3806

36 Crescenzo R, Abate F, Lasorsa E, et al. Convergent mutations and kinase fusions lead to oncogenic STAT3 activation in anaplastic large cell lymphoma. Cancer Cell 2015;27:516–32. doi:10.1016/j.ccell.2015.03.006

37 Vogel TP, Milner JD, Cooper MA. The Ying and Yang of STAT3 in Human Disease. J Clin Immunol 2015;35:615–23. doi:10.1007/s10875-015-0187-8

38 Nairismägi M-L, Tan J, Lim JQ, et al. JAK-STAT and G-protein-coupled receptor signaling pathways are frequently altered in epitheliotropic intestinal T-cell lymphoma. Leukemia 2016;30:1311–9. doi:10.1038/leu.2016.13

39 Sharma A, Oishi N, Boddicker RL, et al. Recurrent STAT3-JAK2 fusions in indolent T-cell lymphoproliferative disorder of the gastrointestinal tract. Blood 2018;131:2262–6. doi:10.1182/blood-2018-01-830968

40 Soderquist CR, Patel N, Murty V V., et al. Genetic and phenotypic characterization of indolent T-cell lymphoproliferative disorders of the gastrointestinal tract. Haematologica 2019;:haematol.2019.230961. doi:10.3324/haematol.2019.230961

41 Jabri B, Sollid LM. Tissue-mediated control of immunopathology in coeliac disease. Nat Rev Immunol 2009;9:858–70. doi:10.1038/nri2670

42 Meresse B, Malamut G, Cerf-Bensussan N. Celiac Disease: An Immunological Jigsaw. Immunity 2012;36:907–19. doi:10.1016/j.immuni.2012.06.006

43 Trynka G, Zhernakova A, Romanos J, et al. Coeliac disease-associated risk variants in TNFAIP3 and REL implicate altered NF-B signalling. Gut 2009;58:1078–83. doi:10.1136/gut.2008.169052

44 O’Keeffe J, Lynch S, Whelan A, et al. Flow cytometric measurement of intracellular migration inhibition factor and tumour necrosis factor alpha in the mucosa of patients with coeliac disease. Clin Exp Immunol 2001;125:376–82. doi:10.1046/j.1365-2249.2001.01594.x

45 Kooy-Winkelaar YMC, Bouwer D, Janssen GMC, et al. CD4 T-cell cytokines synergize to induce proliferation of malignant and nonmalignant innate intraepithelial lymphocytes. Proc Natl Acad Sci 2017;114:E980–9. doi:10.1073/pnas.1620036114

46 Grivennikov SI, Karin M. Dangerous liaisons: STAT3 and NF-κB collaboration and crosstalk in cancer. Cytokine Growth Factor Rev 2010;21:11–9. doi:10.1016/j.cytogfr.2009.11.005

47 Strebovsky J, Walker P, Lang R, et al. Suppressor of cytokine signaling 1 (SOCS1) limits NFκB signaling by decreasing p65 stability within the cell nucleus. FASEB J 2011;25: 863–74. doi:10.1096/fj.10-170597

48 Groner B, von Manstein V. Jak Stat signaling and cancer: Opportunities, benefits and side effects of targeted inhibition. Mol Cell Endocrinol 2017;451:1–14. doi:10.1016/j.mce.2017.05.033

49 Salas A, Hernandez-Rocha C, Duijvestein M, et al. JAK–STAT pathway targeting for the treatment of inflammatory bowel disease. Nat Rev Gastroenterol Hepatol 2020;17:323–37. doi:10.1038/s41575-020-0273-0

50 Nunes AT, Annunziata CM. Proteasome inhibitors: structure and function. Semin Oncol 2017;44:377–80. doi:10.1053/j.seminoncol.2018.01.004

51 Kleppe M, Koche R, Zou L, et al. Dual Targeting of Oncogenic Activation and Inflammatory Signaling Increases Therapeutic Efficacy in Myeloproliferative Neoplasms. Cancer Cell 2018;33:29–43.e7. doi:10.1016/j.ccell.2017.11.009

