## Supplemental_files for "Oncogenetic Landscape Of Lymphomagenesis In Coeliac Disease"

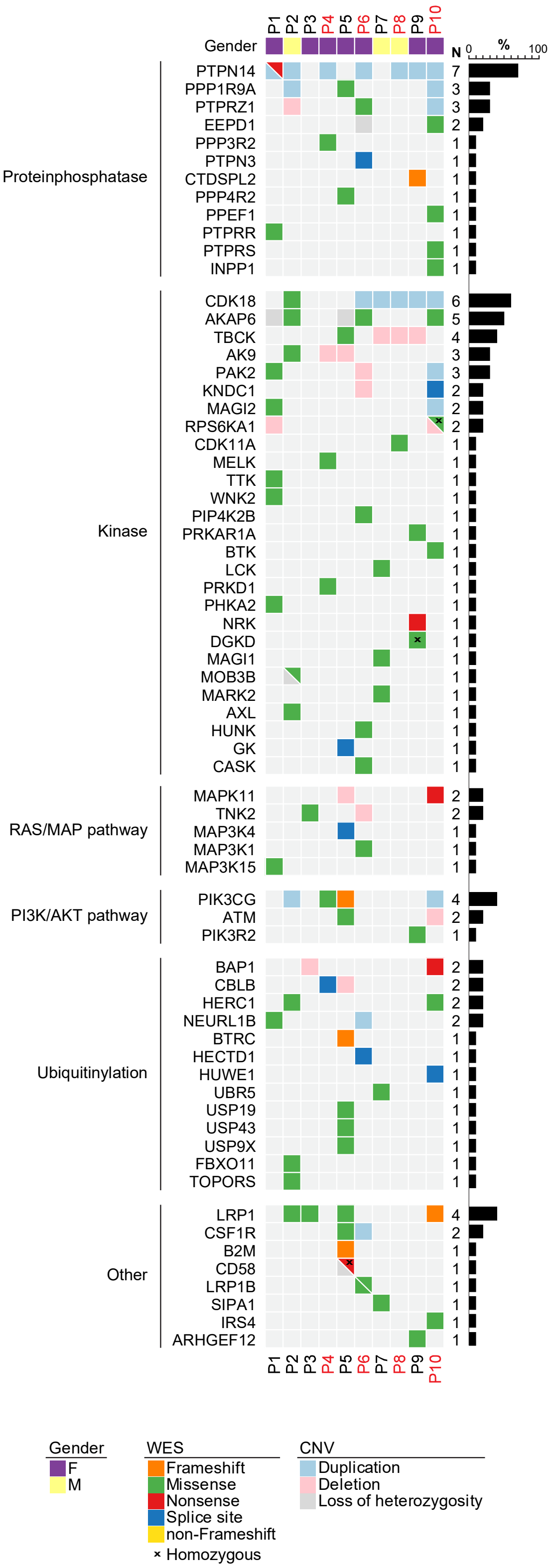


#### Supplementary Figure 1

**To Figure 1: Genomic characterization of the mutational landscape of RCDII cells.** Heatmap summarizes selected somatic variants in combination with CNV results per patient (column) and gene (row), grouped by pathway or function. Mutations are color coded according to the type of aberration as indicated in the legend.

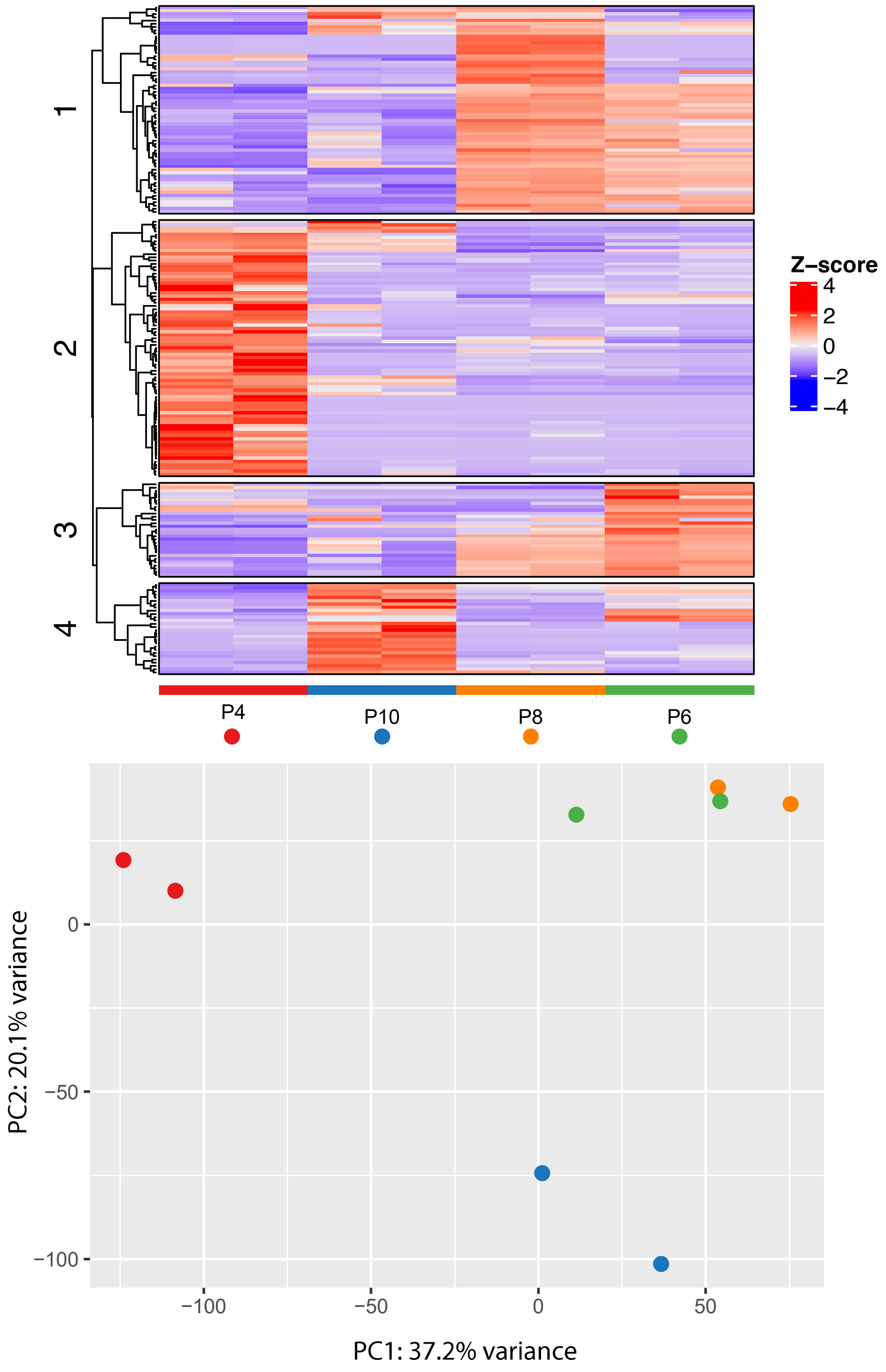


#### Supplementary Figure 2

**To Figure 1:** Individual transcriptional profiles of selected RCDII-cell lines. Upper heatmap shows top 200 differentially expressed genes found in four selected RCDII-cell lines and bottom plot illustrates principal component (PC) analysis with PC1 vs. PC2.

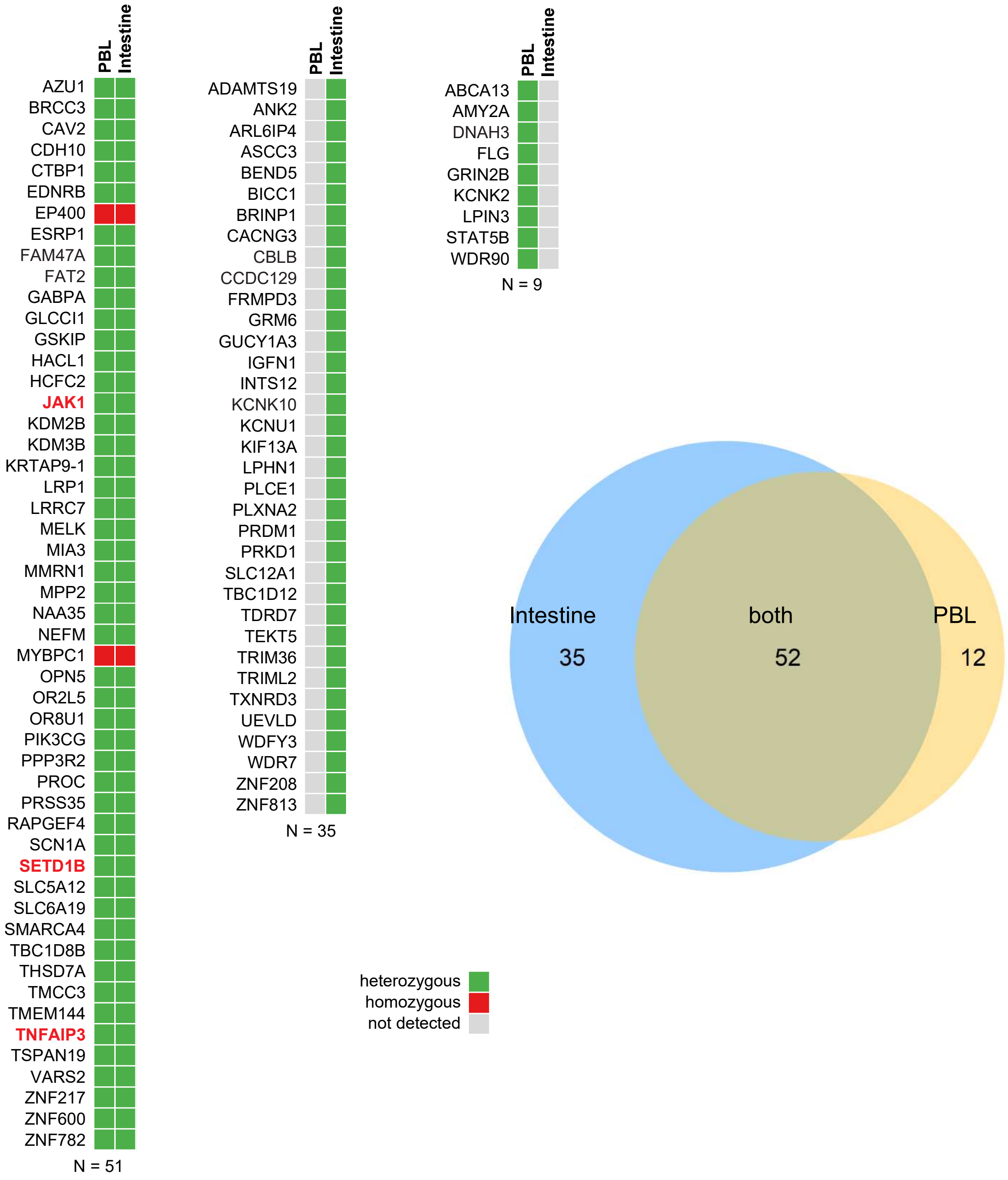


#### Supplementary Figure 3

**To Figure 1:** Individual mutational profiles of RCDII cells derived from intestinal biopsy and freshly isolated from PBL. Venn diagram illustrates overlapping and individual proportion of mutations detected in RCDII-cells derived by culture from duodenal biopsies (Intestine) or directly sorted from peripheral blood (PBL) of the same patient.

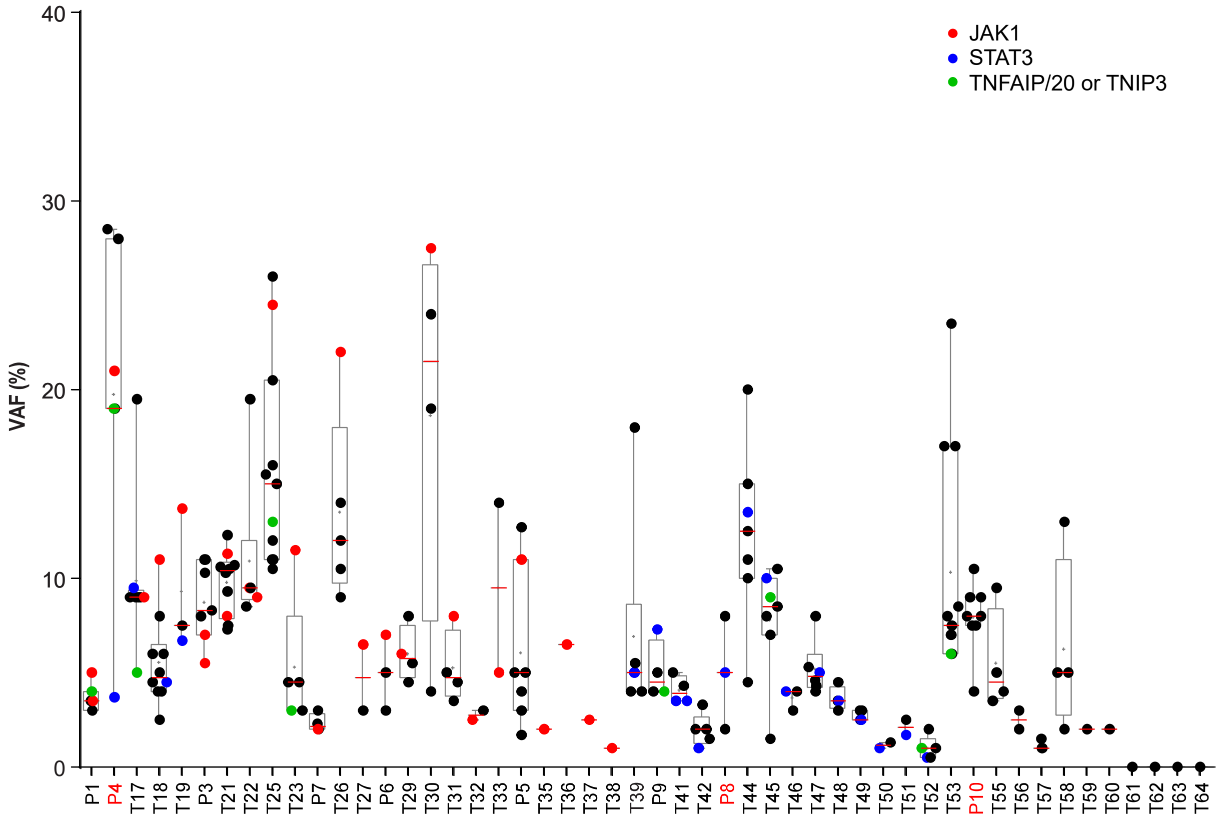


#### Supplementary Figure 4

**To Figure 2: Distribution of variant allele frequencies in RCDII biopsies.** Scatter graph depicts frequency of variant allele frequency (VAF) detected per patient with each dot representing the mean VAF for every single gene called from various runs and techniques. Overlayed box (5-95 percentile) and whiskers ranging from min to max. Red dots indicate JAK1 mutations, blue dots STAT3 mutations and green dots TNFAIP3/A20 or TNIP3.

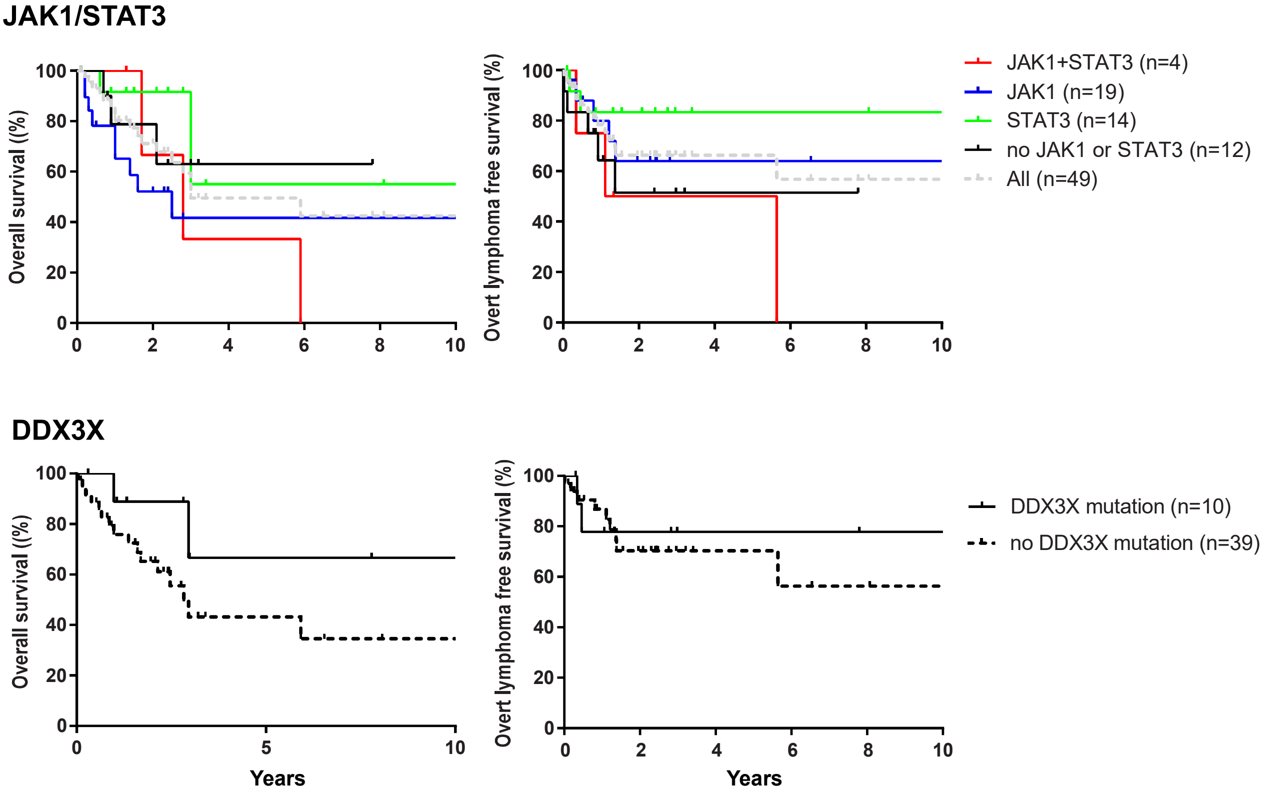


#### Supplementary Figure 5

**To Figure 2. Survival curves.** Overall survival and overt lymphoma free survival curves associated to JAK1/STAT3 and DDX3X mutations.

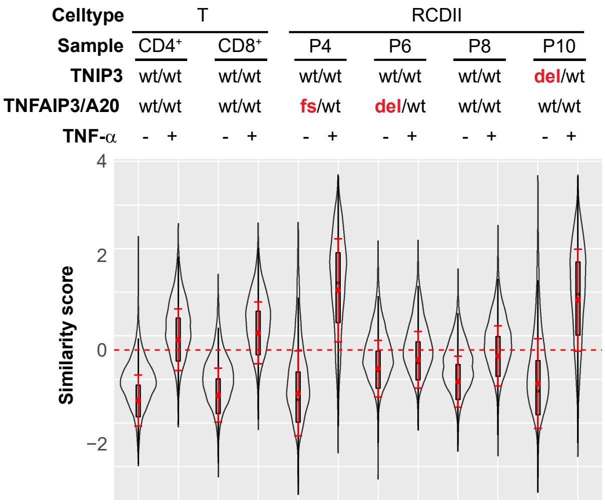


#### Supplementary Figure 6

**To Figure 3: TNFα-induced NFkB activation in RCDII lines.** Violin plot shows translocation scores for NFκB/p50 in non-stimulated or TNFα stimulated RCDII cell lines from four patients and control CD3+CD8+ and CD3+CD4+ T cell lines.

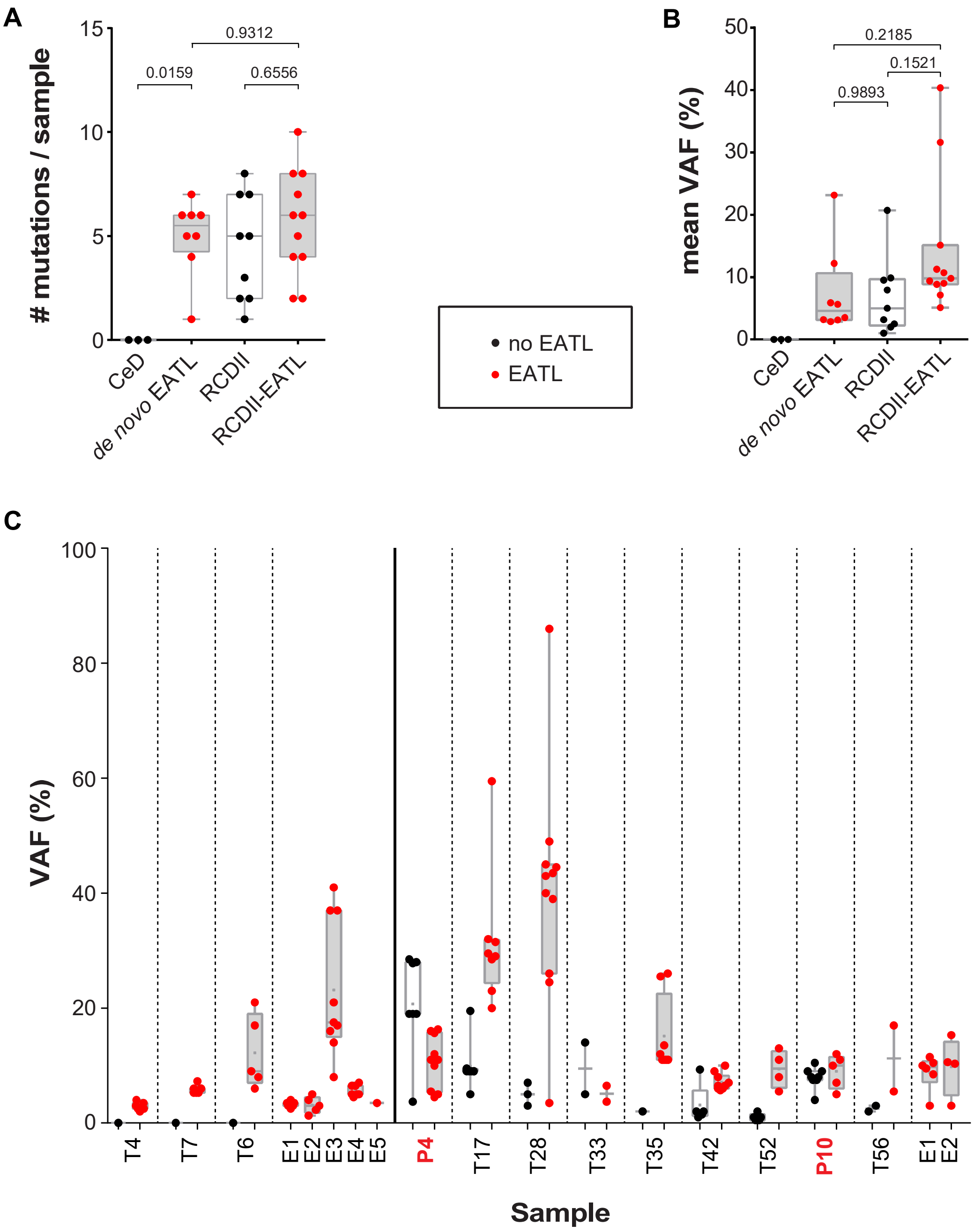


#### Supplementary Figure 7

**To Figure 4:** Distribution of variant allele frequencies in EATL biopsies. (**A**) Box and whisker plots illustrating the number of mutations or (**B**) mean variant allele frequency (VAF) per sample. Indicated p-values were calculated by ANOVA with Tukey correction for multiple comparisons. (**C**) Scatter graph depicts VAF detected per patient with each dot representing the mean VAF for every single gene called from various runs and techniques for *de novo* EATL and RCDII-EATL as indicated. Black dots indicate VAF from autologous samples without EATL (no-EATL) and red dots from EATL samples. Box for 75%ile and whiskers ranging from min to max, horizontal lines are median and grey dots indicating mean.

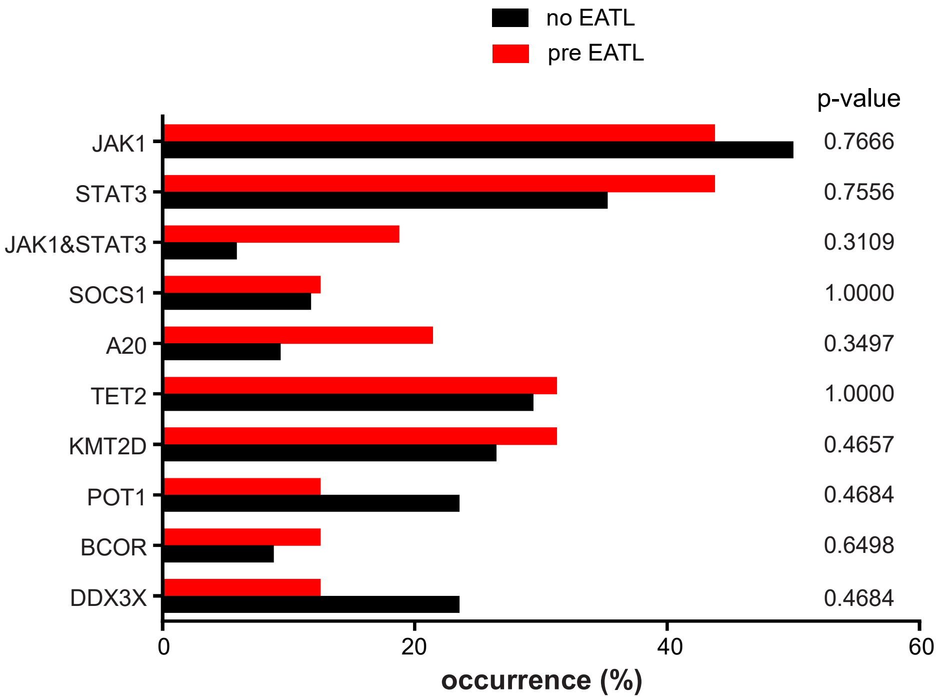


#### Supplementary Figure 8

**To Figure 4:** Predictive value of genes mutated in RCDII for the risk to develop EATL. Bar plot shows occurrence of selected genes which were mutated in biopsies of RCDII patients who developed (red bars = “pre EATL”) or did not develop (black bars = “no EATL”) EATL. Numbers indicate p-values as calculated via Fisher’s exact test.

### SUPPLEMENTARY METHODS

#### Diagnostic procedures

Diagnosis of CeD was based on HLA-DQ2/8 typing, detection of anti-tissue transglutaminase and/or anti-endomysium antibodies and presence of villous atrophy with increased numbers of IEL in intestinal biopsies. Refractory coeliac disease (RCD) was defined by persistent malnutrition syndrome and villous atrophy after one year of strict adherence to GFD and was further divided into type I RCD (RCDI) and RCDII. As reported, RCDI was diagnosed in the absence of detectable monoclonal TCRγ-rearrangement with normal CD8+ T-IEL phenotype. Conversely, RCDII was characterized by monoclonal TCRγ-rearrangement and, in 47/50 patients, by the presence of >50% CD3+CD8- IEL by immunohistochemistry of formalin-fixed paraffin-embedded (FFPE) sections, and/or >25% CD45+CD103+ IEL negative for surface CD3 expression by flow cytometry after isolation from fresh biopsies.[4–7] In 3/50 otherwise typical cases of RCDII, RCDII-IEL consisted of CD8-CD4- IEL that expressed surface TCRγδ (70% and 95% respectively) or TCRαβ (95%). EATL were diagnosed as described.[7–9]

#### Cell culture

Cells were cultured in RPMI (Invitrogen, Thermo Fisher Scientific, Villebon-sur-Yvette, France) supplemented with 10% AB-positive human serum (PAA Laboratories and Sigma-Aldrich, Saint-Quentin Fallavier, France), 1% sodium pyruvate, 1% nonessential amino acids, 1% HEPES buffer, 1μg/ml fungizone, 40μg/ml gentamicin and 5x10–5M β-mercaptoethanol (all Invitrogen) containing 20ng/ml human IL-15 (R&D Systems, Bio-Techne, Lille, France). For STAT3 phosphorylation kinetic studies cells were starved from IL-15 (R&D Systems) in complete

RPMI for 6h, then re-stimulated with 20ng/ml IL-15 (R&D Systems) and harvested at various time points.

**RNA isolation and RNAseq analysis**

Total RNA was isolated using the RNeasy Plus Kit (QIAGEN, Courtaboeuf, France) including a DNAse treatment step. RNA quality was assessed using RNA Screen Tape 6000 Pico LabChips with the Tape Station (Agilent Technologies, Les Ulis, France) and RNA concentrations were measured by spectrophometry using the Xpose (Trinean NV, Gentbrugge, Belgium). Libraries were prepared starting from 1µg of total RNA (for RNA-profiling) or 250ng of total RNA (for fusion gene analysis) using the Universal Plus mRNA-Seq kit (Nugen, Tecan, Lyon, France) as recommended by the manufacturer. The oriented cDNA produced from the poly-A+ fraction was sequenced on a NovaSeq6000 from Illumina (Paired-End reads 100 bases + 100 bases). A total of ~50 million (for expression profiling) or ~100 million (for fusion gene analysis) of passing filter paired-end reads were produced per library. Data was processed by the Bioinformatics core facility at IMAGINE Institute. Fastq raw data was mapped using Hisat2 and gene counts were generated with featureCounts.

**Fusion gene analysis**

Three different tools for gene fusion detection in RNAseq reads were used. STAR-fusion[10] (STAR v2.7.2a, STAR-Fusion v1.7.0 in combination with GRCh38 genome and GENCODE31 annotations), arriba (STAR v2.6.1.d, arriba v1.1.0, GRCh38, GENCODE31; https://github.com/suhrig/arriba/) and fusioncatcher[11] (v1.1.0, GRCh38, annotation human v95) were used to analyze gene fusion events. For the latter, fusions were further prioritized and annotated using oncofuse-1.1.1-1. Candidate fusions that were found by at least two out of the three tools were individually reviewed: 27 gene pairs were consistently identified by at least two of the fusion callers but all but 3 of these were intrachromosomal events between neighboring genes. Upon closer inspection the three putative interchromosomal events were supported by very few reads and did not involve known cancer genes and were therefore discarded.

**T cell receptor rearrangements**

Rearrangements of T cell receptor (TCR) δ,-γ and γ-chains were assessed on genomic DNA from RCDII cell lines and from frozen biopsies by multiplex polymerase chain reaction (PCR) with fluorescent primers according to BIOMED-2 Concerted Action protocols.[1–3]

**Genome-Wide Array-Based Comparative Genomic Hybridization (CGH)**

Array-based CGH (aCGH) was performed on genomic DNA from cultured RCDII-cells. Sex-matched standard reference DNA from pooled human individuals (Agilent Technologies) served as control. Samples were analysed using SurePrint G3 Cancer CGH+SNP (Agilent Technologies). Labelling and hybridization were performed according to the manufacturer's instructions (Agilent Technologies). Array slides were analysed with the Agilent scanner (G2505C) with Feature Extraction software (version 10.1.1.1). Data was analysed with Agilent CytoGenomics software (version 2.0; Agilent Technologies).

**Whole Exome Sequencing (WES)**

Genomic DNA was extracted from leucocytes. Exome capture was performed with the SureSelect Human All Exon kit (Agilent Technologies). Agilent Sure Select Human All Exon (58Mb V6) libraries were prepared from 3 µg of genomic DNA sheared with an Ultrasonicator (Covaris) as recommended by the manufacturer. Barcoded exome libraries were pooled and sequenced with a HiSeq2500 system (Illumina), generating paired-end reads. After demultiplexing, sequences were mapped on the human genome reference (NCBI build 37, hg19 version) with BWA. After demultiplexing, Variant calling was carried out with the Genome

Analysis Toolkit (GATK), SAMtools, and Picard tools. The mean depth of coverage of the exome libraries was greater than ~130X in average with >96 to 99% of the targeted exonic bases covered at least by 15 independent reads and >93 to 98% by at least 30 independent sequencing reads (98-99% at 15X and 93 to 97% at 30X). All the variants were annotated and filtered with PolyWeb, an in-house-developed annotation software.

**Targeted Next Generation Sequencing**

For targeted next-generation sequencing (TNGS) of a custom-made panel of genes involved in lymphomagenesis, genomic DNA-libraries were generated from 50ng of genomic DNA obtained from frozen biopsies and enriched in exonic fragments using the Nextera Rapid Capture method (Illumina, San Diego, CA). All exons encompassing the 104 selected genes (**Supplementary File 1**) were captured with cRNA baits designed with Design Studio (Illumina). Targeted regions were sequenced on a MiSeq or NextSeq system (Illumina). Obtained depth of coverage for each sample was at least 100X.

**Targeted amplicon sequencing**

Mutations identified by TNGS were validated by targeted amplicon sequencing (TAS) using QIAseq Targeted DNA Panels (QIAGEN) designed to target somatically mutated regions from a total of 69 candidate genes previously identified in TNGS and total exons of 22 genes in which potentially oncogenic mutations were identified in WES (**Supplementary File 2**). The starting material consisted of 50ng genomic DNA. The amplified fragments were sequenced as 150bp paired reads using an Illumina NextSeq (Illumina). For reporting, a sequencing coverage of 250X (bidirectional true paired-end sequencing) and a variant frequency of 1% in the wild-type background were used as cut-offs. Sequences produced allowed respectable sequence coverage of 100-25000 reads per bp position, with more than 95% of targeted bases covered.

**Genomic data processing**

WES, TNGS and TAS data were first processed by the bioinformatics platform of Université de Paris. Paired-end sequences were mapped to the GRCh37 (hg19) human genome reference using the Burrows-Wheeler Aligner. Calling of single nucleotide polymorphisms and small indels were performed using Genome Analysis Toolkit (GATK), SAMtools, as well as Picard, as described (http://www.broadinstitute.org/gatk/guide/topic?name = best-practices). Annotation was based on the 72nd version of the ENSEMBL database. Further variant calling was achieved with Freebayes[12] for TNGS and smCounter (https://ngsdataanalysis2.qiagen.com/) for TAS. Sequence data were next visualized using the in-house Polyweb database and variant calls were filtered with PolyQuery (WES) or PolyDiag (TNGS, TAS) interfaces as somatic *de novo* mutations in diseased samples, restricted to impactful coding exonic sequences (exclusion of UTR, silent, intronic, ncRNA, mature miRNA, Pseudogene, synonymous, Up/Downstream, intergenic sequences). Mutations with variant allele frequencies (VAF) of <45% were considered as somatic and VAF at 50±5% as constitutive. For TNGS and TAS, a minimal VAF threshold of 1% was applied. Calls were further filtered for read depth (≥50 total reads, ≥15 alternative reads; ≥10 alternative reads, if variants were located in a hotspot). Sequence quality was individually inspected with Integrative Genomics Viewer.[13] Elimination of irrelevant and frequent polymorphisms was based on frequencies extracted from public databases such as US National Center for Biotechnology Information database of SNP (dbSNP), 1000 Genomes Project, Exome Variant Server (EVS), and Exome Aggregation Consortium (ExAC, http://exac.broadinstitute.org). Candidate mutations were evaluated based on their predicted impact on protein function using three algorithms: Polyphen2 (http://genetics.bwh.harvard.edu/pph2/), SIFT (Sorting Intolerant From Tolerant, J. Craig Venter Institute) and CADD (Combined Annotations Dependent Depletion[14]). Curation of mutations was further achieved through online database repositories such as VarElect (Keywords: Lymphoid, Neoplasm, ALL, LNH) for prioritization, COSMIC (Catalogue of Somatic Mutations in Cancer) Cancer Gene Census and ClinVar for disease association, PeCan (Saint-Jude Pediatric Cancer Data Portal) for hotspot identification, as well as literature research for causal information. Variant rules were defined for 4 classes (definitely pathogenic, probably pathogenic, unknown variant, SNP) according to combinations of above mentioned criteria.

**Flow cytometry**

For staining, $10^5$ cells were incubated for 20 minutes at 4°C with 8-color mixes of FITC-CD103 (2G5, mouse IgG2a, Beckman Coulter), PE-panTCRαβ (IP26A, mouse IgG1, Beckman Coulter), BV510-CD3ε (SK7, mouse IgG1, κ, SONY Biotechnology Inc.), PerCP-Cy5.5-CD45 (2D1, mouse IgG1, κ, SONY Biotechnology Inc), APC-H7 CD8α (SK1, mouse IgG1, κ, BD Biosciences), APC-NKp46 (9E2, mouse IgG1, κ, SONY Biotechnology Inc), V450-CD19 (HIB19, mouse IgG1, κ, BD Biosciences). For intracellular CD3ε staining, cells were fixed and permeabilized using BD Cytofix/Cytoperm kit (BD Biosciences), and labelled with PeCy7-CD3ε (SK7, mouse IgG1, κ, SONY Biotechnology Inc.) stained for 30 minutes at 4°C. Cells were analysed on FACSCanto II using FlowJo software (FlowJo Inc, Ashland, USA). Cell sorting was performed using FACSAriaII (BD Biosciences).

Apoptosis was determined using AnnexinV Apoptosis detection kit (SONY Biotechnology Inc) with propidium iodide and cell proliferation was analysed with Ki-67 (SONY Biotechnology Inc) after 3 days of stimulation. Cells were also stained with anti-human PerCP Cy5.5 CD45 (2D1, mouse IgG1, κ, SONY Biotechnology Inc), anti-human BV510 CD3ε (SK7, mouse IgG1, κ, SONY Biotechnology Inc). Dead cells were excluded with 7-ADD (SONY Biotechnology Inc.) for apoptosis detection and Life/Dead Near IR (Life Technologies) and AnnexinV-APC (SONY Biotechnology Inc) for Ki-67 staining. Data were acquired on LSR Fortessa or FACSCantoII (Both BD Biosciences) flow cytometer and analysed with FlowJo version 10 software.

**Western blot**

Cells were harvested and subsequently lysed in RIPA lysis buffer with protease inhibitor (cOmplete Mini™, Roche) and phosphatase inhibitor cocktails 2 and 3 (Sigma-Aldrich). Proteins were separated under denaturizing conditions in 8%-12% SDS-PAGE gels (Bio-Rad Laboratories, Marnes-la-Coquette, France) and transferred to PVDF membranes (Bio-Rad Laboratories). The following primary antibodies were used: pSTAT3 (3E2; mouse monoclonal), STAT3 (124H6; mouse monoclonal) and β-actin (AC-15; HRP-conjugated mouse mAb, Sigma). Secondary antibody was anti-mouse IgG (goat) and blots were revealed with Clarity and Clarity Max ECL (Bio-Rad Laboratories). All antibodies were from Cell Signaling Technology (Ozyme, Saint Quentin-en-Yvelines, France) if not otherwise stated.

**Imaging flow cytometry**

For NFκB translocation analysis, $1x10^6$ cells were stimulated with 100ng/ml recombinant TNFα (BioLegend, Ozyme) for 20 minutes, subsequently subjected to the Amnis NFκB Translocation kit (Luminex Corp., Austin, USA) according to the manufacturer's instructions with 7-AAD as nuclear counterstain. Image acquisition was performed at 60X magnification with the ImageStream XMkII multispectral imaging flow cytometer (Amnis Corp., Seattle, USA), and acquired images were analyzed with the IDEAS software (version 6.2; Amnis Corp.). Cytoplasmic to nuclear translocation of NFκB/p50 transcription factor was measured using the similarity score, which quantifies the intensity values of the nuclear and cytoplasmic NFκB protein image pixels.

**Drug Inhibition assays**

CD103+sCD3– RCDII-cell lines were derived from duodenal biopsies of RCDII patients and maintained as previously described[15,16]. For drug testing, RCDII cell lines were cultured at $10^6$ cells/mL in medium supplemented with 20ng/mL IL-15 (R&D Biosystems) in presence of 20nM ruxolitinib (Sellekchem, Euromedex, Souffelweyersheim, France), 0,01µM and 1µM abrocitinib (PF-04965842; Sellekchem), 1µM and 20µM budesonide (Sigma Aldrich), 10nM and 100nM bortezomib (Sellekchem) or dimethylsulfoxyde (Sigma Aldrich) as control. Inhibition of growth and induction of apoptosis were assessed by flow cytometry using the cell cycle marker KI67 and AnnexinV and propidium iodide respectively, while inhibition of STAT3 phosphorylation was assessed by Western blot.

**Data visualization**

WES and CGH data were visualized with circos-0.69-6.[17] Graphs were generated with Prism 6 software (GraphPad Software Inc., San Diego, USA) or R-3.6.2 and RStudio with ggplot2-3.3.1[18] (PCA, violin plot), ComplexHeatmap-2.2.0[19] (heatmap), VennDiagram-1.6.20[20] (venn diagram) or corrplot-1.0.0[21] (co-occurrence).

REFERENCES

1 Ettersperger J, Montcuquet N, Malamut G, *et al.* Interleukin-15-Dependent T-Cell-like Innate Intraepithelial Lymphocytes Develop in the Intestine and Transform into Lymphomas in Celiac Disease. *Immunity* 2016;**45**:610–25. doi:10.1016/j.immuni.2016.07.018

2 van Dongen JJM, Langerak AW, Brüggemann M, *et al.* Design and standardization of PCR primers and protocols for detection of clonal immunoglobulin and T-cell receptor gene recombinations in suspect lymphoproliferations: Report of the BIOMED-2 Concerted Action BMH4-CT98-3936. *Leukemia* 2003;**17**:2257–317. doi:10.1038/sj.leu.2403202

3 Derrieux C, Trinquand A, Bruneau J, *et al.* A Single-Tube, EuroClonality-Inspired, TRG Clonality Multiplex PCR Aids Management of Patients with Enteropathic Diseases, including from Formaldehyde-Fixed, Paraffin-Embedded Tissues. *J Mol Diagnostics* 2019;**21**:111–22. doi:10.1016/j.jmoldx.2018.08.006

4 Cellier C, Patey N, Mauvieux L, *et al.* Abnormal intestinal intraepithelial lymphocytes in refractory sprue. *Gastroenterology* 1998;**114**:471–81. doi:10.1016/S0016-5085(98)70530-X

5 Malamut G, Afchain P, Verkarre V, *et al.* Presentation and Long-Term Follow-up of Refractory Celiac Disease: Comparison of Type I With Type II. *Gastroenterology* 2009;**136**:81–90. doi:10.1053/j.gastro.2008.09.069

6 Rubio-Tapia A, Murray JA. Classification and management of refractory coeliac disease. *Gut* 2010;**59**:547–57. doi:10.1136/gut.2009.195131

7 Cheminant M, Bruneau J, Malamut G, *et al.* NKp46 is a diagnostic biomarker and may be a therapeutic target in gastrointestinal T-cell lymphoproliferative diseases: A CELAC study. *Gut* 2018;**68**:1396–405. doi:10.1136/gutjnl-2018-317371

8 Swerdlow SH, Campo E, Harris NL, *et al.* Summary for Policymakers. In: Intergovernmental Panel on Climate Change, ed. *Climate Change 2013 - The Physical Science Basis*. Cambridge: : Cambridge University Press 2017. 1–30. doi:10.1017/CBO9781107415324.004

9 Malamut G, Verkarre V, Callens C, *et al.* Enteropathy-Associated T-Cell Lymphoma Complicating an Autoimmune Enteropathy. *Gastroenterology* 2012;**142**:726-729.e3. doi:10.1053/j.gastro.2011.12.040

10 Haas BJ, Dobin A, Li B, *et al.* Accuracy assessment of fusion transcript detection via read-mapping and de novo fusion transcript assembly-based methods. *Genome Biol* 2019;**20**:213. doi:10.1186/s13059-019-1842-9

11 Nicorici D, Satalan M, Edgren H, *et al.* FusionCatcher - a tool for finding somatic fusion genes in paired-end RNA-sequencing data. 2014. doi:10.1101/011650

12 Garrison E, Marth G. Haplotype-based variant detection from short-read sequencing -- Free bayes -- Variant Calling -- Longranger. *arXiv Prepr arXiv12073907* Published Online First: 2012. doi:arXiv:1207.3907 [q-bio.GN]

13 Robinson JT, Thorvaldsdóttir H, Winckler W, *et al.* Integrative genomics viewer. *Nat Biotechnol* 2011;**29**:24–6. doi:10.1038/nbt.1754

14 Kircher M, Witten DM, Jain P, *et al.* A general framework for estimating the relative pathogenicity of human genetic variants. *Nat Genet* 2014;**46**:310–5. doi:10.1038/ng.2892

15 Mention J-J, Ben Ahmed M, Bègue B, *et al.* Interleukin 15: a key to disrupted intraepithelial lymphocyte homeostasis and lymphomagenesis in celiac disease. *Gastroenterology* 2003;**125**:730–45. doi:10.1016/S0016-5085(03)01047-3

16 Malamut G, El Machhour R, Montcuquet N, *et al.* IL-15 triggers an antiapoptotic pathway in human intraepithelial lymphocytes that is a potential new target in celiac disease-associated inflammation and lymphomagenesis. *J Clin Invest* 2010;**120**:2131–43. doi:10.1172/JCI41344

17 Krzywinski M, Schein J, Birol I, *et al.* Circos: An information aesthetic for comparative genomics. *Genome Res* 2009;**19**:1639–45. doi:10.1101/gr.092759.109

18 Wickham H. *ggplot2*. New York, NY: : Springer New York 2009. doi:10.1007/978-0-387-98141-3

19 Gu Z, Eils R, Schlesner M. Complex heatmaps reveal patterns and correlations in multidimensional genomic data. *Bioinformatics* 2016;**32**:2847–9. doi:10.1093/bioinformatics/btw313

20 Chen H, Boutros PC. VennDiagram: A package for the generation of highly-customizable Venn and Euler diagrams in R. *BMC Bioinformatics* 2011;**12**:35. doi:10.1186/1471-2105-12-35

21 Wei T, Simko V, Levy M, *et al.* Visualization of a Correlation Matrix. *Statistician* Published Online First: 2017.https://github.com/taiyun/corrplot

**Supplementary Table 1: Characteristics of EATL patients**

| | **Total (N=19)** | ***de novo* EATL (N=8)** | **RCDII-EATL (N=11)** | **Stat** |
|---|---|---|---|---|
| **Clinical features** | | | | |
| **Age**, mean in years (range) | 55.9 (38-73) | 57.4 (38-73) | 55 (37-71) | ns |
| **Gender** (F, %) | 12 (63%) | 4 (50%) | 8 (73%) | ns |
| **Time from CD diagnosis and molecular evaluation,** mean in years (range) | 4.7 (0-21.7) | 1.7 (0-10.9) | 6.8 (0.6-21.7) | **p=0.0064** |
| **Biological features** | | | | |
| **TCR status** (frozen biopsies) | | | | |
| Gamma clonal | 19 (100%) | 8 (100%) | 11 (100%) | ns |
| VDJ Beta clonal | 4/18 (22%) | 3/7 (43%) | 1 (9%) | ns |
| **Mutational status** (frozen biopsies) *JAK1/STAT3* mutations | 15 (79%) | 8 (100%) | 7 (63%) | ns |
| *JAK1* | 14 (74%) | 8 (100%) | 6 (55%) | **p=0.045** |
| *STAT3* | 10 (50%) | 2 (25%) | 7 (64%) | ns |
| *TNFAIP3* mutations | 5/18 (28%) | 0 | 5/10 (50%) | **p=0.036** |
| *TNIP3* mutations | 1/18 (6%) | 1 (13%) | 0/10 | ns |
| *DDX3X* mutations | 6 (32%) | 2 (25%) | 4 (36%) | Ns |

**Supplementary Table 2: Characteristics of RCDII patients according to *JAK1* and *STAT3* status**

| | **Total (N=50)** | ***JAK1*** | | | ***STAT3*** | | |
|---|---|---|---|---|---|---|---|
| | | **Mutated (N=24)** | **Not mutated (N=26)** | **Stat** | **Mutated (N=19)** | **Not mutated (N=31)** | **Stat** |
| **Clinical features** | | | | | | | |
| **Age**, mean in years (range) | 57 (29-77) | 57 (30-77) | 57 (29-77) | ns | 54 (37-71) | 59 (29-77) | ns |
| **Gender** (F, %) | 30 (60%) | 13 (54%) | 17 (65%) | ns | 14 (74%) | 16 (52%) | ns |
| **Time from CeD diagnosis to molecular evaluation,** mean in years (range) | 5.4 (0.1-21.9) | 5.3 (2.7-17.9) | 5.4 (0.1-21.9) | ns | 4.5 (0.1-10.9) | 5.9 (0.2-21.9) | ns |
| **History of EATL** (%) | 16 (32%) | 8 (33%) | 8 (31%) | ns | 6 (32%) | 10 (32%) | ns |
| **Biological features** | | | | | | | |
| **Flow cytometry** (N=), small bowel biopsies | 33 | 15 | 18 | | 14 | 19 | |
| % aberrant population in IEL (/CD45$^+$) | 72% (23-98) | 80% (46-98) | 65% (23-89) | **p=0.04** | 71% (23-95) | 73% (30-98) | ns |
| % aberrant population in LPL (/CD45$^+$) | 32% (5-70) | 43% (6-94) | 23% (5-70) | **p=0.03** | 30% (6-60) | 34% (5-70) | ns |
| % aberrant population in PBL (/CD45$^{hi}$) | 7% (0-80) | 12% (0-80) | 3% (0-11) | ns | 11% (0-80) | 4% (0-25) | ns |
| **TCR status** (frozen biopsies) | | | | | | | |
| Gamma clonal | 48 (96%)[#] | 23 (96%)[#] | 25 (96%)[#] | ns | 19 (100%) | 29 (94%)[#] | ns |
| VDJ Beta clonal | 15/49 (31%) | 9/23 (39%) | 6/26 (23%) | ns | 6/19 (32%) | 9/30 (30%) | ns |
| **Mutational status** (frozen biopsies) | | | | | | | |
| *JAK1* mutations | 24 (48%) | - | - | | 5 (26%) | 19 (61%) | **p=0.02** |
| *STAT3* mutations | 19 (38%) | 5 (21%) | 14 (54%) | **p=0.02** | - | - | - |
| *TNFAIP3/TNIP3* mutations | 10/47 (21%) | 4/21 (19%) | 6 (23%) | ns | 7/18 (39%) | 3/28 (11%) | **p=0.03** |
| *DDX3X* mutations | 10/50 (20%) | 4 (21%) | 6 (23%) | - | 4 (21%) | 6 (19%) | ns |

[#] two cases with polyclonal TCRγ and clonal TCRδ

**Supplementary Table 3: Characteristics of RCDII patients according to *DDX3X* and *TNFAIP3/TNIP3* status**

| | **Total** (N=50) | ***DDX3X*** | | | ***TNFAIP3/TNIP3*** | | |
|---|---|---|---|---|---|---|---|
| | | **Mutated** (N=10) | **Not mutated** (N=40) | **Stat** | **Mutated** (N=10) | **Not mutated** (N=36) | **Stat** |
| **Clinical features** | | | | | | | |
| **Age**, mean in years (range) | 57 (29-77) | 56 (29-71) | 57 (30-77) | ns | 53 (34-77) | 59 (30-77) | ns |
| **Gender** (F, %) | 30 (60%) | 4 (40%) | 25 (63%) | ns | 5 (50%) | 24 (67%) | ns |
| **Time from CeD diagnosis to molecular evaluation,** mean in years (range) | 5.4 (0.1-21.9) | 5.8 (2.7-10.9) | 5.3 (0.1-21.9) | ns | 5.3 (0.2-21.7) | 5.4 (0.1-21.9) | ns |
| **History of EATL** (%) | 16 (32%) | 2 (22%) | 14 (35%) | ns | 3 (30%) | 10 (28%) | ns |
| **Biological features** | | | | | | | |
| **Flow cytometry** (N=), small bowel biopsies | 33 | 7 | 26 | | 8 | 24 | |
| % aberrant population in IEL (/$CD45^+$) | 72% (23-98) | 79% (40-95) | 70% (23-98) | ns | 84% (40-98) | 70% (23-98) | ns |
| % aberrant population in LPL (/$CD45^+$) | 32% (5-70) | 36% (8-70) | 31% (5-94) | ns | 49% (8-94) | 28% (5-70) | p=0.06 |
| % aberrant population in PBL (/$CD45^{hi}$) | 7% (0-80) | 6% (0-11) | 7% (0-80) | ns | 19% (0-80) | 2% (0-16) | **p=0.02** |
| **TCR status** (frozen biopsies) | | | | | | | |
| Gamma clonal | 48/50 (96%)[#] | 10/10 (100%) | 38/40 (95%)[#] | ns | 10 (100%) | 34 (94%)[#] | ns |
| VDJ Beta clonal | 15/49 (31%) | 3/10 (30%) | 12/39 (31%) | ns | 1 (10%) | 12 (33%) | ns |
| **Mutational status** (frozen biopsies) | | | | | | | |
| *JAK1/STAT3* mutations | 38 (74%) | 7 (70%) | 31 (78%) | ns | 9 (90%) | 25 (69%) | ns |
| *JAK1* | 24 (48%) | 4 (40%) | 20 (50%) | ns | 4 (40%) | 17 (47%) | p=0.07 |
| *STAT3* | 19 (38%) | 4 (40%) | 15 (38%) | ns | 7 (70%) | 12 (34%) | P=0.06 |
| *TNFAIP3/TNIP3* mutations | 10/47 (21%) | 2/9 (22%) | 8/38 (21%) | ns | - | - | - |
| *DDX3X* mutations | 10/50 (20%) | - | - | - | 2 (20%) | 6 (17%) | ns |

[#] two cases with polyclonal TCRγ and clonal for TCRδ
